# Interconnections with and within the trypanosomal respiratory chain revealed by complexome profiling

**DOI:** 10.64898/2026.09.08.750046

**Authors:** Corinna Benz, Rayyan Tariq Khan, Michael Hammond, Ondřej Gahura, Ingrid Škodová-Sveraková, Urban Koblar, Lawrence Rudy Cadena, Philip Thomas Butterill, Marek Vrbacký, Hassan Hashimi

**Author notes:** **Correspondence:** Hassan Hashimi, Institute of Parasitology, Biology Centre, CAS, v.v.i., Branisovska 31, 370 05 Ceske Budejovice, Czech Republic, E-mail address. Current addresses: Johannes Kepler University Linz and Kepler University Hospital GmbH, Department of Pathology and Molecular Pathology, Linz, Austria (Urban Koblar). Current address: Department of Biology, Institute of Microbial Cell Biology, Heinrich Heine University Düsseldorf, Düsseldorf, Germany (Lawrence Rudy Cadena).

## Abstract

Proteins are frequently integrated into multicomponent complexes that execute the elaborate processes supporting life. Thus, a protein’s function can only be defined by the company it keeps within a complex. Proteomes provide informative protein inventories but lack information about protein quaternary structures. Complexome profiling (CP) has been transformative in capturing the comprehensive population structure of complexes at a given moment within a cell. We have employed CP to chart the multiprotein complex landscape of two life cycle stages of *Trypanosoma brucei.* This data has allowed the observation of previously hidden interactions with and within the mitochondrial respiratory chain. We have found (1) an exceptional case of a SLC25 solute transporter that interacts with NADH dehydrogenase, (2) two ATP synthase subunit g paralogs that are intriguingly excluded from the enzyme’s dimers, and (3) refined the known composition of ubiquinol:cytochrome c oxidoreductase by addition of missing subunits and removing an incorrectly assigned subunit, which more likely acts to insert the iron-sulfur co-factor into the complex. Further investigation into ubiquinol:cytochrome c oxidoreductase assembly revealed crosstalk between incorporation of its nuclear subunits with mitochondrial translation, possibly facilitating a hitherto unknown quality control mechanism. These discoveries demonstrate the power of our CP data for generation and testing of hypotheses concerning the multiprotein complexes of the mitochondrion and other organelles of *T. brucei*, a protist of medical and evolutionary importance.

**Statement for broader audience:** Mitochondria are proverbial ‘cellular powerhouses’ thanks to enclosing the respiratory chain (RC), a collection of multiprotein complexes. Their complete make-up and interactions beyond the RC remain unresolved. We have cataloged the proteins of RC and other mitochondrial complexes in unicellular Trypanosoma, which remodels its RC during its life cycle. We reveal and investigate interesting interconnections uncovered by our ‘complexome’ to expand the known interactions that the RC engages in to keep the power running in mitochondria.

## Introduction

Multiprotein complexes carry out the majority of processes essential for cellular viability (Wang *et al*, 2009). This statement certainly applies to the complexes that comprise the mitochondrial respiratory chain: NADH dehydrogenase (Complex I), succinate dehydrogenase (Complex II), ubiquinol:cytochrome c oxidoreductase (Complex III), cytochrome c oxidase (Complex IV) and F_O_F_1_-ATP synthase (Complex V). (Ryan *et al*, 2013). Stability of subunits in a complex is often interdependent (Ryan *et al*., 2013) and their mutations often exhibit the same phenotype as a result of critical disrupted function (Lage *et al*, 2007).

Another common feature of protein complexes is the recruitment of lineage-specific subunits around a core of evolutionarily conserved proteins (Prokopchuk *et al*, 2023). Consequently, while the existence of some complexes can be inferred from the presence of genes encoding conserved subunits, their complete composition remains unknown due to the incorporation of novel subunits or subunits whose primary structure has diverged beyond recognition by homology searches. Filling these knowledge gaps is not just a matter of bookkeeping as these unaccounted peripheral subunits impact the protein complexes that house them. Such additions can lead to changes in the processes that the affected complex executes (*e.g.* Kaurov *et al* (2018)). For instance, the respiratory chain complexes are especially prone to accumulating microproteins, which are less than 100 amino acids long (Hassel *et al*, 2023), which pose another obstacle for homology searches. This is especially true for the microprotein subunits that have become indispensable building blocks of Complex I (Shin *et al*, 2025). Less frequently but very consequential, is the fact that even core subunits of ancient complexes can become diversified beyond recognition by conventional homology searches (Sheikh *et al*, 2025).

The importance of better understanding the complete composition of individual protein complexes motivated systems wide (*e.g.* Acestor *et al* (2011)) examination of purified protein complexes by basic proteomic approaches, such as liquid chromatography-tandem mass spectrometry. While such studies have advanced our understanding of the complexes residing in various cell types, there are some drawbacks to this approach. Among them are the reliance on transgenesis to introduce handles to capture complexes and the implementation of purification pipelines susceptible to detection of false positive and/or negative interactions. Furthermore, *a priori* knowledge is needed to select a bait to purify a putative complex, with no guarantee that the affinity-handle will not disrupt native and/or introduce spurious interactions.

Complexome profiling (Cabrera-Orefice *et al*, 2022) circumvents the shortcomings of targeted complex purification. Native proteins are separated by their molecular weight via assorted methods, commonly blue native polyacrylamide gel electrophoresis (BN-PAGE). Proteins with similar distribution profiles within an array of size fractions are clustered computationally. Thus, within an assayed cell or organelle, the population structure of multiprotein complexes is ascertained, *i.e.* the configuration and abundance of all detected complexes at once. Complexome profiling can place an orphan protein into its native complex (Heide *et al*, 2012), map stepwise assembly intermediates (Guerrero-Castillo *et al*, 2017), discover mitochondrial protein import quality control mechanisms (Schulte *et al*, 2023), reveal how membrane phospholipid defects affect complex assembly (Van Strien *et al*, 2019), and map the protein complex landscape in apicomplexan parasites (Maclean *et al*, 2021), including changes during life-cycle progression (Evers *et al*, 2021).

Here, we map the mitochondrial complexomes of two life cycle stages of the kinetoplastid protist, *Trypanosoma brucei*. During its life cycle alternating between the tsetse fly vector and mammalian host, oxidative phosphorylation (OXPHOS) is respectively active and suppressed (Bílý *et al*, 2021; Zíková, 2022). The procyclic form (PCF) that occupies the insect midgut bears a canonical mitochondrion that produces ATP by OXPHOS. In the bloodstream form (BSF), however, Complexes III and IV are not expressed. Instead, cell division is sustained only by glycolysis, a metabolic pathway that is mostly encapsulated in modified peroxisomes called glycosomes (Quiñones *et al*, 2020).

We acknowledge a prior complexome profiling study performed on PCF *T. brucei* (Gazestani et al, 2016), which has been employed to refine the interaction network among the subunits of the complex that performs the core enzymatic activities of U-insertion/deletion RNA editing (Poorinmohammad & Salavati, 2024). In the previous study, native complexes were separated by density gradient ultracentrifugation and ion exchange chromatography, biochemical methods, which resulted in an underrepresentation of membrane associated proteins (Gazestani et al., 2016). As we are particularly interested in integral membrane complexes, which construct mitochondria ultrastructure (Colina-Tenorio *et al*, 2026; Pánek *et al*, 2020), we employed BN-PAGE, which better preserves such complexes (Wittig *et al*, 2006). As a consequence, none of the novel interacting proteins we report here were assigned to complexes in the previous study. During revision of this article, another gel-based trypanosomatid complexome study was published (Solayman *et al*, 2026); we provide an in-depth comparison of our complexome data to theirs below.

Fortunately, efforts to eliminate the human disease caused by *T. brucei* have been successful, although completion of this goal remains challenging (Barrett *et al*, 2024). Nevertheless, *T. brucei* remains a relevant laboratory model thanks to transgenic tools that can test hypotheses sparked by omics data, such as those from complexome profiling. Not only can *T. brucei* inform us on the biology of kinetoplastids, a group that still contains causative agents of various diseases globally, but they also have the potential to inform us about eukaryotic biology in general (Ginger *et al*, 2026). The last common ancestor of eukaryotes was likely an excavate, a taxon now represented by an assemblage of flagellated protists on either side of the root of the eukaryotic tree of life (Williamson *et al*, 2025). For now, *T. brucei* is the most powerful lab model from this evolutionarily important group (Hashimi, 2019), and therefore has utility for illuminating the epoch before the diversification of extant eukaryotes.

## Results and Discussion

### The complexome of two life cycle stages of Trypanosoma brucei

To investigate the composition of the mitochondrial protein complexes that inhabit metabolically different life cycle stages, we performed complexome profiling of enriched PCF and BSF mitochondria. After premeabilization by the mild non-ionic detergent digitonin, we separated the complexes as a function of their respective sizes using two types of BN PAGE gels with different polyacrylamide gradients. Each was optimized to better resolve either larger (3-12%) or smaller (4-16%) molecular weight complexes (*e.g.* Fig. 1B). Prior to the mass spectrometry phase of the CP pipeline, we confirmed these conditions were appropriate for downstream complexome analysis by immunoblotting which showed the expected three major Complex V bands corresponding to F_0_F_1_ ATP synthase dimers, followed by monomers and a soluble F_1_ moiety (Fig. S1A).

**Figure 1.**
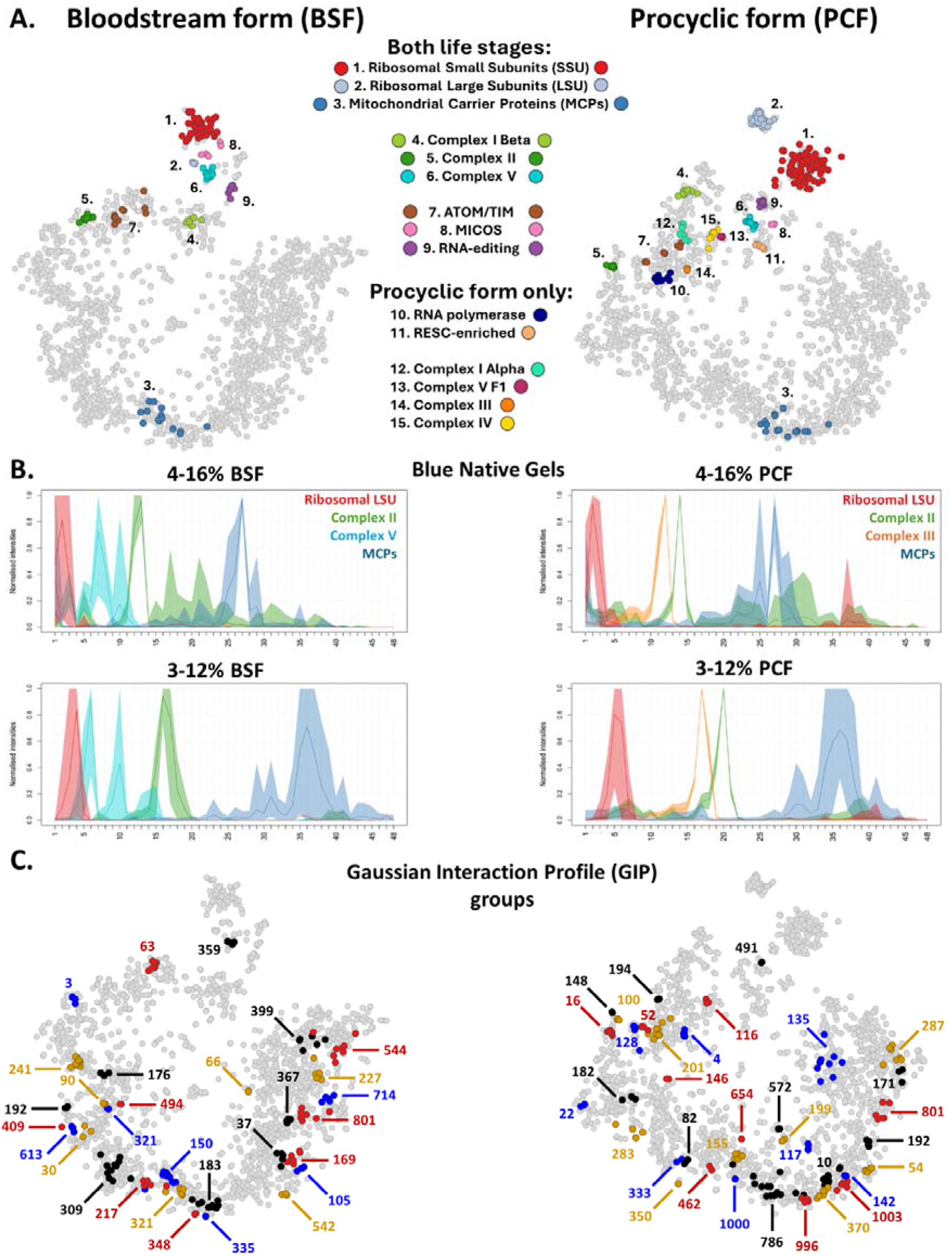
Combined complexome profiles of bloodstream and procyclic form *Trypanosoma brucei*. **(A)** The distribution of subunits of marker complexes (color-coded as indicated in the central key) in combined complexome profiles (*i.e.* BSF∪ and PCF∪) visualized as a t-SNE plot. **(B)** Size fraction distribution of individual complexome profiles color coded as indicated on top right of upper graphs. See key in A for abbreviations. x-axis, molecular weight size fraction 1 (largest) to 48 (smallest); y-axis, normalized signal intensity. **(C)** Fractionally distinct Gaussian Interaction Profile (GIP) groups from combined complexome profiles plotted onto t-SNE plot in A. Group identification number of each group as listed in Table S1. See Figure S2 for t-SNE plots of complexome profiles from individual gels.

Next, we employed these conditions for the complexome analysis. Organelles enriched from PCF and BSF were solubilized and resolved by BN-PAGE, and each lane was cut into 48 slices. The distribution of relative abundance of all proteins throughout 48 gel slices was calculated by intensity Based Absolute Quantitation (iBAQ) (Cabrera-Orefice *et al*., 2022). Mitochondrial and glycosomal proteins were well represented in our datasets (Dataset S1), with the former being more abundant in PCF as expected (Table 1).

**Table 1.** Number of detected proteins and predicted Gaussian Interaction Profiles (GIPs) in each complexome dataset. Last 2 rows contain statistics from Solayman and colleagues (2026). N/A, not applicable; *L. tar., Leishmania tarentolae*.

|  |  |  | **Proteins detected** |  |  | **GIP totals** |  |  |
|---|---|---|---|---|---|---|---|---|
| **Dataset** | **Stage** | **Gel** | **Total** | **Mitochondrial<sup>1</sup>** | **Glycosome<sup>2</sup>** | **Cluster #** | **Protein #** | **Ave protein/GIP** |
| PLK139 | BSF | 3-12% | 2414 | 685 | 61 | 75 | 289 | 3.9 |
| PLK107 | BSF | 4-16% | 3492 | 607 | 74 | 165 | 719 | 4.4 |
| N/A | BSF | Combined | N/A | 500 | 57 | 58 | 304 | 5.2 |
| PLK139 | PCF | 3-12% | 2526 | 1005 | 62 | 92 | 398 | 4.3 |
| PLK107 | PCF | 4-16% | 3616 | 1034 | 79 | 174 | 718 | 4.1 |
| N/A | PCF | Combined | N/A | 906 | 61 | 69 | 358 | 5.2 |
| Solayman | BSF | N/A | 2094 | 1027 | 37 | 45 | 167 | 3.7 |
| Solayman | PCF | N/A | 2126 | 1228 | 48 | 139 | 633 | 4.6 |
| Solayman | *L.tar.* | N/A | 2790 | 1238 | N/A | 128 | 642 | 5.0 |

**<sup>1</sup>Based on (Vacas *et al.*, 2026); <sup>2</sup>based on (Güther *et al.*, 2014)**

Histograms of Complex V F_1_ subunit protein intensity correlate with α-ATPase across immunoblot resolutions in each gel (Fig. S1B). Furthermore, F_o_ protein intensity shows co-accumulation in immunoblot bands corresponding to its F_o-_F_1_ dimer, while being absent in the F_1_ moiety (Fig. S1A,B), consistent with the known architecture of the complex (Gahura *et al*, 2022b). We additionally note a slight accumulation of F_o_ intensity in the lower molecular weight regions of PCF gels, but not those of BSF (Fig. S1B), as might be reasonably expected, given the higher relative abundance of Complex V in PCF. In conclusion, there is correspondence between complex profiles of individual Complex V subunits and immunoblot detection of the complex.

To further validate the complexome datasets, we assessed the presumed clustering of known mitochondrial complex subunits belonging to the mitoribosome, respiratory chain complexes, MICOS, RNA polymerases, and RNA editing complexes. We resolved the combined abundance across gel slices of PCF and BSF *via* t-distributed stochastic neighbor embedding (t-SNE), showing individual proteins clustered in two-dimensional space based on the similarity of their fractionation profiles. This was performed for all four individual datasets (Fig. S2A-B) and additionally through a combination of both PCF (PCF∪) and BSF (BSF∪) complexome datasets (Fig. 1A). Such resolution demonstrated that the majority of these complex subunits exhibited fractional co-similarity. In certain cases, fractional differences allowed complexes to be split into sub-clusters, such as the mitoribosome small (mtSSU) and large subunits (mtLSU), as well as the subunits of Complex I alpha and beta subcomplexes (Acestor *et al*., 2011; Surve *et al*, 2012). Variance among life stages was also observed, with the F_o_ and F_1_ subunits of complex V only resolved separately in PCF, likely due to their aforementioned lower molecular weight accumulation (Fig. S1B) while Complexes III, IV and alpha subunits of Complex I were absent or diminished in BSF (Fig. 1A). MICOS is assembled in both life cycle stages despite the lack of Complexes III and IV in the BSF, as reported previously (Boudová *et al*, 2026). A minority of known complex subunits fractionated differently than the rest of the complex, showing enrichment in lower size fractions, most likely due to their dissociation from the respective complex under our lysis and separation conditions. However, our data demonstrate that complex integrity has overall been preserved, as observed by co-fractionation.

We additionally noted that the majority of detected mitochondrial carrier proteins (MCPs), belonging to the SLC25 family of solute carriers (Kunji *et al*, 2025), showed similar distribution profiles (Fig. 2A). However, this is more likely influenced by the similar mature protein molecular weights (43-78 kDa) than forming a traditional complex. MCP4 represents a notable deviation (Tb927.10.4910), but this can be explained by its outlier size of 78 kDa (Colasante *et al*, 2009). Of particular note is one MCP which consistently associates with Complex I, which we discuss below.

**Figure 2.**
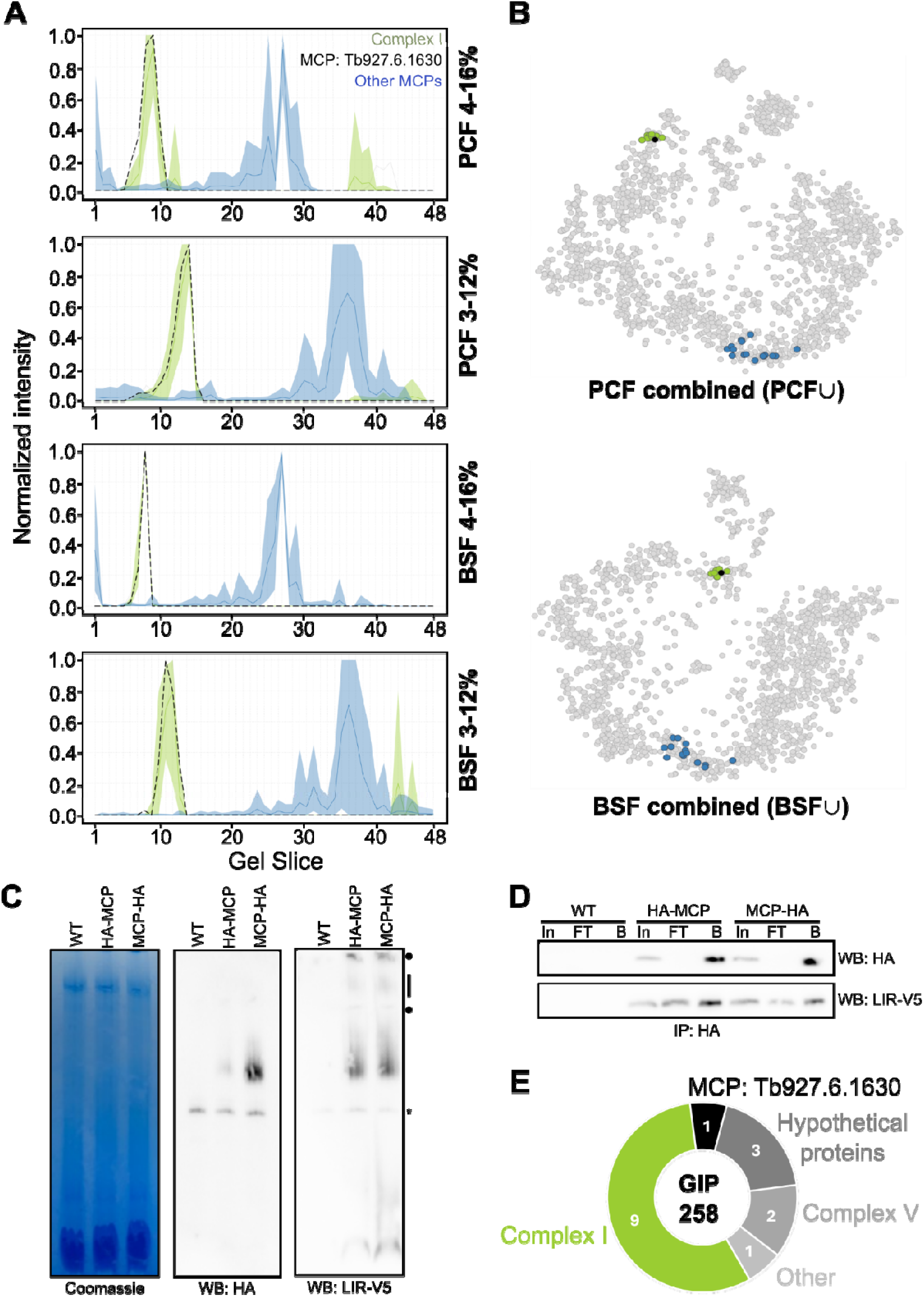
A mitochondrial carrier protein interacts with the Complex I beta subcomplex. **(A)** The relative abundancies of conventional mitochondrial carrier proteins (MCPs, blue lines) and MCP Tb927.6.1630 (dashed black line), which co-migrates with Complex I beta (green line), across the 48 size fractions in procyclic form cells. As in Fig. 1B. Color coding key on top right of top plot. **(B)** t-SNE plots from BSF∪ and PCF∪ showing distribution of proteins in A using same color code as in A. **(C)** BN PAGE detecting the MCP tagged on the N-(HA-MCP) and C-(MCP-HA) termini and V5-tagged Complex I beta subunit LIR. Samples were solubilized in 2% Dodecyl Maltoside (DDM). Circles and line shown on right demark higher molecular weight species of LIR-V5 that do not co-migrate with the MCP. Antibody used to probe western blots (WB) below each image. Wild type (WT) cells as a control for antibody specificity; *, non-specific band, running at approximately 720 kDa. **(D)** Western blots (antibody indicated on right) showing immunoprecipitation (IP) of HA-MCP and MCP-HA (Top) and co-IP with V5-tagged LIR (Bottom). In, input; FT, flow thru and B, beads (immunocaptured protein) with WT serving as a negative control. **(E)** Pie chart showing the composition of GIP group 258 from 4-16% BSF complexome profile dataset. See Table S1 for composition of GIP group.

### A mitochondrial carrier protein interacts with Complex I

We noticed in all four profiles that a 39 kDa MCP (Tb927.6.1630) migrated at ∼1 MDa in a position corresponding to considerably larger complexes than the other MCPs (Fig. 2A). Upon further examination, this MCP clustered with Complex I beta subunits (Surve *et al*., 2012) in both life cycle stages (Fig. 2A-B). To verify this result, we created PCF cell lines in which the Complex I beta subunit LIR (Tb927.11.15440) was appended with the V5 epitope tag and the MCP with either a C- or N-terminal HA tag. Regardless on which terminus the HA epitope was placed, it co-migrated with the lowest molecular weight species of LIR-V5 on BN PAGE (Fig. 2C). This robust association led us to hypothesize that this MCP interacts with Complex I beta. To test this, we performed immunoprecipitation on the MCP tagged on either end. Indeed, LIR-V5 co-immunoprecipitated with both versions of the tagged MCP (Fig. 2D), confirming the interaction.

It is important to note that a minor portion of LIR-V5 was not immobilized in the beads, consistent with the higher molecular weight fractions of LIR-V5 that did not appear to co-migrate with HA-tagged MCP in BN PAGE (Fig. 2D). Together, this suggests that this MCP interacts with most but not all of the LIR-V5 proteins in the cell. This seems to disagree with our complexome data, in which this MCP co-migrates with all Complex I beta (Fig. 2A). This can be explained by the different detergents used in the BN PAGE performed to generate the complexome data and used to observe the migration of the epitope-tagged proteins in Fig. 2C. Equally, LIR may not represent a Complex I subunit directly interacting with this MCP. Of course, the tags themselves may also influence BN PAGE mobilities of the examined proteins compared to the endogenous forms seen in the complexome datasets.

The current paradigm is that MCPs function as monomers, with some members of this family forming homodimers (Kunji *et al*., 2025). Thus, their interaction with other classes of proteins is rather exceptional. For example, the MCP responsible for ATP/ADP antiport across the inner membrane has been reported to interact with the TIM23 inner membrane protein translocase (Mehnert *et al*, 2014) and Complex III and IV (Dienhart & Stuart, 2008) in yeast, plus F_O_F_1_-ATP synthase in rats (Chen *et al*, 2004). However, this is not a universal phenomenon as the ATP/ADP carrier does not interact with these respiratory chain complexes in *T. brucei* (Gnipová *et al*, 2015). Here, our complexome revealed the unexpected interaction between an MCP and Complex I. As far as we know, this is the first instance of an MCP other than the ATP/ADP carrier to be shown to interact with a respiratory chain complex. This situation is likely restricted to kinetoplastids and may be connected to *T. brucei* Complex I losing the capacity to generate proton motive force and being split into two discrete subcomplexes (Surve *et al*., 2012). Perhaps, our finding may help to shed light on the enigmatic function of Complex I *e.g.* by determining which solute(s), if any, is translocated by this MCP. In a broader context, this finding may help to explain how certain adaptations are employed as Complex I becomes reduced, which has occurred independently in assorted eukaryotic lineages (Glastad & Johnston, 2025).

### Unbiased clustering to predict protein complexes by Gaussian mixture modeling

To interrogate our complexome data beyond known protein complexes, we employed this dataset to predict novel protein assemblies. Proteins with similar complexome profiles were clustered using the Gaussian Interaction Profiler (GIP), which implements a Gaussian mixture model and bootstrapping to confidently assign subpopulations within a dataset (van Strien *et al*, 2024). We note that more GIPs were assigned in individual PCF versus BSF datasets, and a lower amount of GIPs were assigned when the two complexome profiles from each stage were combined (Fig. 1C, Table 1). The average number of proteins assigned per GIP was ∼4 in the individual versus ∼5 in the combined datasets (Fig. S3A; Table S1). All the GIPs exhibit similar size fraction dispersion, with most GIPs found between fractions 15 and 38 (Fig. S3D; Table S1), approximately representing ∼900 to ∼40 KDa (Fig. S3B).

GIP designation can be observed correlating to certain established mitochondrial complexes. For example, among BSF∪ GIP 359 contains a selection of complex V subunits, while GIP 150 is enriched for a number of MCPs (c.f. Fig. 1A and C, Table S1). We also observed that the Complex I-associating MCP was reconstructed through gaussian mixture modelling with Complex I in GIP group 258 from the 4-16% BSF complexome profile dataset (Fig 2E). Accordingly, while this Complex I-MCP fractional similarity can be observed across all datasets (Fig 2. A,B), the association is strongest in 4-16% BSF, which thus represents the optimized analysis conditions to view this relationship. Among PCF∪, GIP 116 contains numerous subunits of Complex IV, while GIP 16 showed a similar distribution for subunits of Complex II (Fig. 1A&C encircled, Table S1).

GIP group 4 from the PCF∪ dataset appeared to correspond to Complex III of the respiratory chain (Fig. 1A&C encircled,, S4A). This group contains 10 proteins, half of which are known *T. brucei* Complex III subunits (Acestor *et al*., 2011). In contrast, only three commonly recognizable Complex III subunits (Table S2, Group 49 TbCF-HC net) were clustered together in the previous complexome profile study in which membrane proteins were underrepresented (Gazestani *et al*., 2016). In our combined PCF dataset, 4 hypothetical proteins were found and corroborated through their shared presence in GIP groups generated against the individual BN PAGE complexome profiles. We dubbed these UQCRTBx1-4 (Table S2) to signify these are putative *T. brucei* subunits and decided to investigate them further to verify the veracity of our complexome data.

### Divergent ubiquinol:cytochrome c oxidoreductase subunits are predicted by Gaussian mixture modeling

We decided to test whether the four UQCRTBx1-4 proteins are indeed subunits of Complex III as predicted by GIP (Fig 3A, S4A). First, we asked whether any of them are homologous to mitochondrial proteins that were identified in the solved structure of Complex III from *Euglena gracilis* (He *et al*, 2024), encapsulated with *T. brucei* in the taxon Euglenozoa (Kostygov *et al*, 2021). However, reciprocal homology searches (Table S2) only identified five of the previously identified Complex III subunits (Acestor *et al*., 2011) plus a diverged ortholog of the Hinge protein UQCRH. While UQCRH was placed in the Complex III GIP group 618 in the 3-12% PCF dataset, it was ultimately excluded from the UQCRTBx candidate proteins as it did not cluster with Complex III in both PCF complexome profiles. FoldSeek structural homology searches (van Kempen *et al*, 2024) were also attempted to determine the identity of UQCRTBx1-4 and confirm that of UQCRH (Table S1). Only UQCRH returned a moderately confident hit, while UQCRTBx1-4 did not. Interestingly, UQCRTBx4 did return a very weak hit to mouse UQCR10.

**Figure 3.**
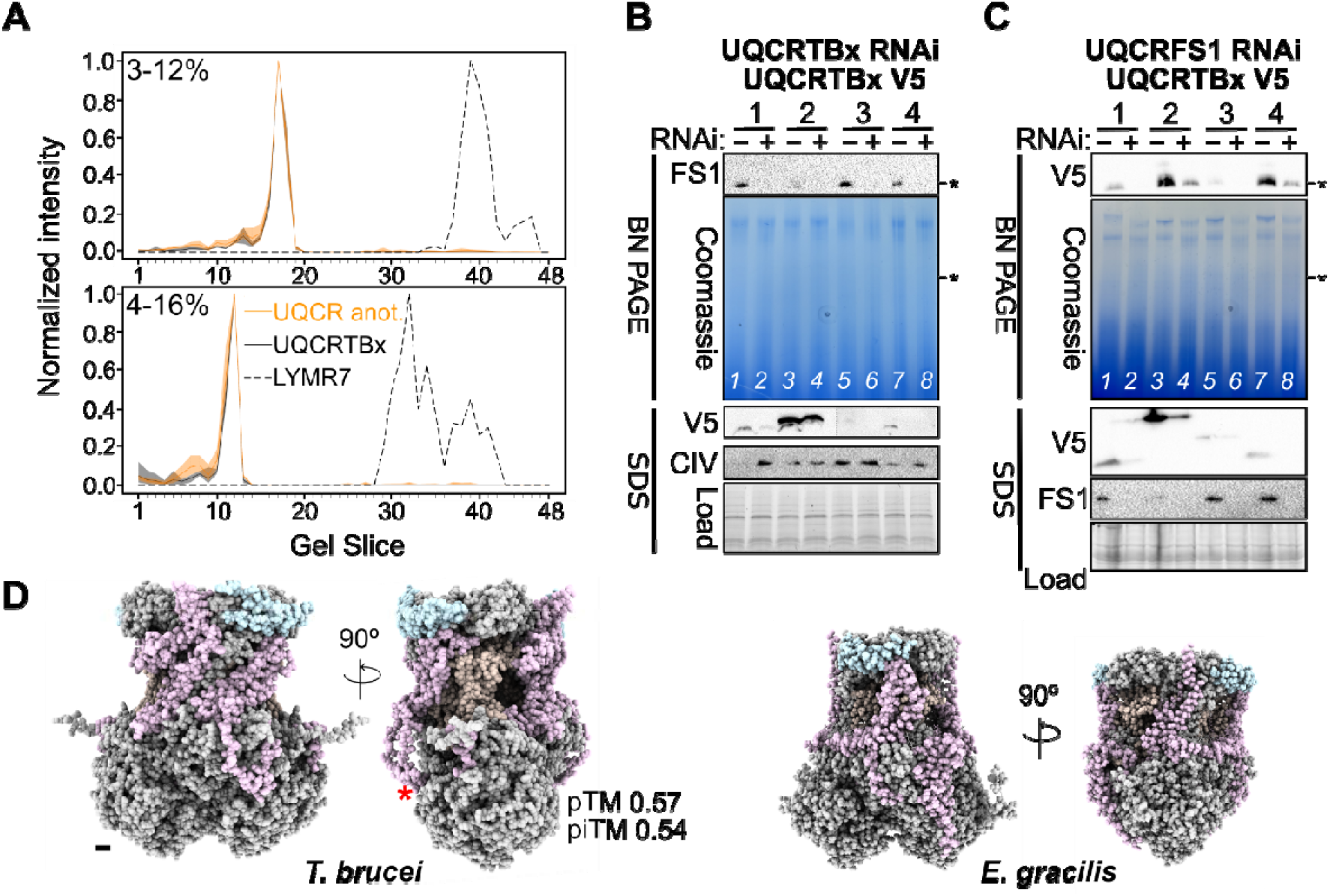
Identification of four divergent components of ubiquinol: cytochrome c oxidoreductase. **(A)** The relative abundancies of known Complex III subunits (solid black line), a potential interactor, LYRM (dashed black line) and the four divergent subunits (yellow line) across the 48 size fractions in procyclic form cells. As in Figure 1B. **(B)** Top: BN PAGE (BN) and western blot detecting Rieske protein (FS1) in induced (+) vs non-induced (-) UQCRTBx1-4 RNAi cell lines. RNAi was induced for four days. Samples solubilized in 1.5% w/v digitonin (Dig.). Bottom: SDS PAGE (SDS) and western blot of the same samples detecting V5-tagged TBx1-4 and Complex IV (CIV) component CoxIV. Part of the stain-free gel is shown as a loading control. Asterisks denote 720 kDa marker. **(C)** Top: BN PAGE and western blot detecting V5-tagged UQCRTBx1-4 in induced (+) vs non-induced (-) Rieske protein (FS1) RNAi cell lines. RNAi was induced for four days. Samples solubilized in 1.5% w/v digitonin (Dig.). Bottom: SDS PAGE and western blot of whole cell lysates prepared at the same time as the samples for BN PAGE. V5-tagged TBx1-4 and Rieske protein (FS1) were detected, part of the stain-free gel is shown as a loading control. Asterisks as in B. **(D)** AlphaFold3 prediction of *T. brucei* Complex III containing the four divergent subunits (purple) in comparison to the solved structure of *Euglena gracilis* Complex III (PDB 8IUF) (He *et al*., 2024). UQCRCH identified in one GIP group and by structural homology in light blue. *, location of missing UQCRB subunit in predicted structure. *E. gracilis* orthologs of these proteins colored the same. Scale bar at the bottom left of leftmost model, 10 Å. Related to Figure S3 and Table S2.

As homology searches failed to convincingly assign UQCRTBx1-4 as Complex III subunits, we generated cell lines enabling inducible downregulation and antibody detection of each candidate. These cell lines were verified by immunoblot detection of the V5 epitope appended to the C-terminus of each Complex III subunit candidate, whose signal was reduced after 4 days of RNAi-induction (Fig. 3B). Depletion of each candidate resulted in slower growth (Fig. S4B), which was also observed when known Complex III subunits were downregulated by RNAi (Horváth *et al*, 2005). We hypothesized that the observed slower growth was due to destabilization of Complex III. Indeed, this was observed upon RNAi-silencing of each candidate, as demonstrated by the loss of the ∼720 KDa signal from the Rieske antibody on BN PAGE immunoblots (Fig. 3B). These knockdowns did not influence the levels of Complex IV, diminishing the possibility that this phenotype was unspecific.

We also tested the inverse scenario, in which the stability of the UQCRTBx1-4 proteins depends on the presence of Rieske (also known as UQCRFS1). To reject the null hypothesis, we V5 epitope tagged the candidates in cell lines capable of inducible Rieske downregulation. We observed that each protein was also incorporated into a ∼720 KDa complex, whose assembly was reduced upon Rieske depletion (Fig. 3C). Interestingly, UQCTTBx2-V5 seemed to destabilize the complex prior to RNAi-induction (Fig. 3B, lane 3); this will be discussed in depth in the next section. We conclude from these two experiments that UQCRTBx1-4 are *bona fide* Complex III subunits.

### Divergent ubiquinol:cytochrome c oxidoreductase subunits complete the predicted structure of the complex

We asked how UQCRTBx1-4 would fit with the other known subunits to form the Complex III dimer, the presumable steady-state form of the complex. We entered the protein sequences of each known Complex III subunits plus the four candidates into the AlphaFold3 (AF3) server (Abramson *et al*, 2024). To comply with AF3 size constraints, only one copy of the UBQRB subunit was used, allowing two copies of the rest to be modeled. This predicted a structure that approached the solved *E. gracilis* Complex III (Fig. 3D). Furthermore, the pTM score of the predicted complex is just above the 0.5 threshold considered to approximate the true structure (Rennie & Oliver, 2025). This score is likely reduced by the presence of UQCRTBx1-4, which represent the lowest confidence predicted structures within the structure as per their overall pLDDT scores among (Fig. S5A). They may have also contributed to the ipTM score just below the 0.6 threshold for possible interactions among the amino acids within the structure. Nevertheless, 45 confident interactions were predicted (PDE ≥0.7), with ∼73% of them predicted with high confidence (PDE ≥0.9). Most high confidence contacts are represented by binary interactions between either group of two core proteins UQCRC1 and UQCRC2 or cytochrome b (MT-CYB) and cytochrome c1 (CYC1).

Among the 4 subunits discovered here, UQCRTBx4 and UQCRTBx2 respectively are predicted to make two and one high confidence contacts with the core protein CYC1. Because *E. gracilis* CYC1 is in proximity to UQCR9 (He *et al*., 2024), we wondered if either UQCRTBx4 or 2 may represent the trypanosomal orthologs. By structural alignment, it seems that UQCRTBx4 and 2 may be the UQCR9 and UQCR10 orthologs, respectively (Fig. S5B). The orthology of the former is consistent with the weak FoldSeek hit to mouse UQCR10, which is an ortholog of *E. gracilis* UQCR9 (He *et al*., 2024).

UQCRTBx1 and 3 do not appear to contact other Complex III subunits, which may be a consequence of their low pLDDT scores (Fig. S5A). Nevertheless, UQCRB, which was co-purified with UQCRC1 (Acestor *et al*., 2011), also shows no intermolecular contacts, suggesting that not all actual contacts were confidently predicted by AF3. Structural alignment suggests that UQCRTBx1 and 3 are orthologs of *E. gracilis* UQCREG1 and UQCRQ, respectively (Fig. S5C). The orthology of UQCREG1, a subunit restricted to euglenazoans (He *et al*., 2024) was particularly convincing, showing similar overlap as the UQCRH (Table S1), whose alignment was performed to demonstrate the validity of this distant similarity approach. The localization of UQRCH to the intermembrane space (IMS) via import by oxidative folding is also supported by its depletion when the IMS import pathway is ablated (Kaurov *et al*, 2022; Kaurov *et al*., 2018) (Table S1). Interestingly, UQCRTBx4 is similarly affected, which may fit the proposed model that this protein works with UQRCH to form a CYC1 subassembly during Complex III biogenesis (Geldon *et al*, 2021).

We had observed that introducing a 69 amino-acid long C-terminal V5 tag to UQCRTBx2 by modifying one of the *uqcrtbX2* alleles (Fig. S4C) had a perceptible dominant negative effect on *T. brucei* doubling time compared to the other uninduced cell lines (Fig. S4D). This growth retardation is likely due to potential steric effect of the V5-tag on Complex III assembly and/or stability (Fig. 3B, lane 3). The addition of one copy of UQCRTBx2-V5 to the otherwise same cohort of proteins used for AF3 prediction Complex III (Fig. S5D-E) resulted in a predicted structure with similar pTM and ipTM scores, suggesting that the predicted complexes with and without UQCRTBx2-V5 were equally probable. However, it’s inclusion resulted in much less intra-complex interactions than wild-type Complex III: 35 with ≥0.7 PDE and 27 with ≥0.9 PDE (Fig. S5D), perhaps indicative of an intrinsically unstable complex. Strikingly, contacts between CYC1, UQCRTBx2 and 4 were completely abolished. The predicted structure incorporating UQCRTBx2-V5 is looser, with a more voluminous cavity likely corresponding to one of the quinone (Q) binding sites (Fig S4E). This also suggests that the ability of the altered complex to complete the Q cycle is impaired. Furthermore, the V5 tag occludes the cytochrome c binding site on the IMS face of the predicted complex physically and electrostatically via a positively charged patch introduced onto the electronegative surface. Thus, insertion of UQCRTBx2-V5 could debilitate the complex’s electron transfer activity, necessitating its removal by proteolysis (Deshwal *et al*, 2020), thus explaining the observed reduction in its steady-state level.

Thanks to the discovery of the 4 divergent UQCRTBx subunits via complexome, we are now able to predict the structure of Complex III (Fig. 3D). We have preliminarily designated these subunits based on structural homology to their *E. gracilis* counterparts in the solved Complex III of the euglenid (Fig. S5C; Table S1). After our predictions based on structural homology were made, they were verified in the solved structure of Complex III in *T. brucei* (Hu *et al*, 2026), validating this application of our complexome data.

### A LYRM7 ortholog is likely a ubiquinol:cytochrome c oxidoreductase assembly factor that intermittently interacts with the complex

Interestingly, a ∼20 kDa protein found in the initial purification of Complex III by Acestor and colleagues (2011) did not co-migrate with any of the known and new Complex III subunits in our complexome data (Fig. 3A). FoldSeek–a considerably more sensitive homology search than available at the time of the protein’s initial detection–revealed that this protein is a likely ortholog of LYRM7 (Table S2), a protein responsible for insertion of an iron-sulfur cluster co-factor into the Rieske protein UQCRFS1 (Fernandez-Vizarra & Zeviani, 2018). The protein is named after a conserved LYR motif toward the N-terminus that is important for its function (Dohnálek & Doležal, 2024)(Fig. S4E). A functional link between this putative LYRM7 and Complex III is also suggested by their concomitant upregulation during differentiation from BSF or PCF (Dejung *et al*, 2016).

To gain empirical insight into the putative *T. brucei* ortholog of LYRM7, we epitope tagged the protein. Most of LYRM7-HA migrated at a considerably lower molecular weight than Complex III (Fig. 4A, lane 2), consistent with our complexome data. Interestingly, we observed that a fraction of LYRM7-HA co-migrates with a ∼720 kDa complex. This immunopositive band is indeed Complex III as it disappeared–with a concurrent accumulation of LYRM7-HA to smaller molecular weight bands–when complex assembly was ablated by UQCRTBx1 RNAi (Fig 4A, lane 3). However, the steady state levels of tagged LYRM7 remain the same when UQCRTBx1 or UQCRFS are depleted (Fig 4B). Taken together, it appears LYRM7 interacts intermittently with Complex III. We speculate that the introduction of the HA tag may extend LYRM7 docking onto Complex III in comparison to its endogenous form, which was only detected in lower molecular weight fractions in our complexome datasets.

**Figure 4.**
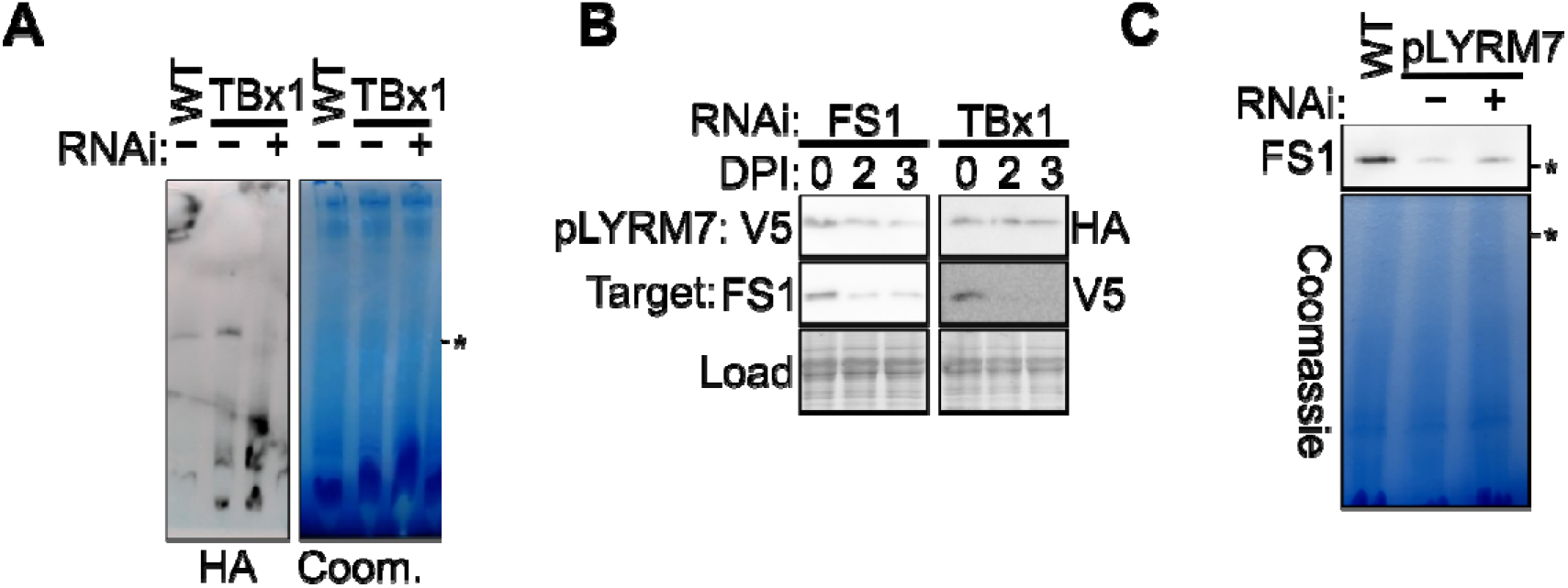
A likely LYRM7 ortholog plays a role ubiquinol: cytochrome c oxidoreductase biogenesis. **(A)** BN PAGE (right) and western blot (left) detecting HA-tagged LYRM7. The Coomassie-stained gel is shown for loading. The asterisk marks 720 kDa. **(B)** Western blot detecting V5- or HA-tagged LYRM7 in Rieske protein (FS1) or UQCRTBx1 (TBx1) RNAi background. The RNAi was induced for zero, two, and three days and verified by also detecting Rieske protein (FS1) or V5 (V5-tagged UQCRTBx1). Part of the stain-free gel is shown as a loading control. **(C)** BN PAGE and western blot detecting FS1 in LYRM7 RNAi induced (+) or non-induced (-) for four days. The Coomassie-stained gel is shown for loading. Asterisks denote 720 kDa marker.

Next, we wanted to test whether LYRM7 depletion affects Complex III assembly. Induction of RNAi against LYRM7 did not affect growth compared to the non-induced control cells (Fig. S4F, G). However, we did notice that the doubling time of the non-induced cells was already increased prior to RNAi-induction, which did not further affect growth drastically (Fig. S4D). We speculate that this observation may be due to leaky RNAi induction in this cell line, a common occurrence in *T. brucei* (Wickstead *et al*, 2002) or the V5 tag interfering with the function of LYRM7, as was seen in epitope tagged UQCRTBx2 (Fig. S4D). Indeed, the non-induced LYRM7 RNAi cell line grows slower than the non-induced cell lines in which one of the four UQCRTBx proteins we characterized were targeted (Fig. S4D). Thus, we conclude that LYRM7 depletion does affect growth, similarly to UQCRTBx1-4 RNAi.

We decided to investigate Complex III assembly in the LYRM7-silenced cell lines in comparison to the parental cell line, *e.g.* wild type PCF (Fig. 4C). We observed that while Complex III was assembled in the LYRM7 cell line regardless of RNAi induction, it was to a considerably lesser extent than in wild type PCF. This phenotype was also observed when the baker’s yeast ortholog of *LYRM7* was deleted (Atkinson *et al*, 2010; Atkinson *et al*, 2011). This was accompanied by a slight but perceptible decrease in steady state Rieske protein during RNAi (Fig. S4G), which we speculate would be more apparent if the non-induced cell line was already not negatively impacted in the first place. Nevertheless, this phenotype is not as severe as the loss of Rieske in *lyrm7* gene deletion yeast (Atkinson *et al*., 2010; Atkinson *et al*., 2011). This discrepancy can be due to technical reasons, as we compare a *lyrm7* gene knockout to the ‘leaky’ RNAi we observed in *T. brucei*. Alternatively, we may have uncovered a *bona fide* a difference in how LYRM7 affects Rieske protein stability in these two lineages.

Because our results were not fully consistent with findings from yeast we closely examined high-confidence AF3 predicted models of yeast and *T. brucei* (Wheeler, 2021) LYRM7 (Fig. S4H) to provide further evidence of their orthology. Both predicted structures have the same three α-helix bundle that contains the LYR motif as well as a downstream two-stranded β-sheet. The latter is not a universal featureof the LYRM superfamily (Dohnálek & Doležal, 2024), making this compelling evidence for our assignation. Furthermore, both examined proteins also have an intrinsically disordered region (IDR) at the C-terminus. The most conspicuous motif that is present in *T. brucei* pLYRM7 but absent from the yeast ortholog is a α-helix pair in between the β-sheets and IDR, which is highly conserved in kinetoplastids (Fig. S4E). Thus, our current working hypothesis is that we have found a genuine LYMR7 ortholog in trypanosomatids, which has been elusive (Dohnálek & Doležal, 2024). Our claim is supported by structural homology (Fig. S4H; Table S2), its apparently transient association with Complex III (Fig. 4A), and a decrease of assembled Complex III upon LYRM7 downregulation (Fig. 4C), which phenocopies *lyrm7* deletion mutants in yeast (Atkinson *et al*., 2010; Atkinson *et al*., 2011).

### An unexpected link between Ubiquinol:cytochrome c oxidoreductase assembly and mitochondrial translation

After refining the definitive subunit composition of *T. brucei* Complex III, we next asked in which order the UQCRTBx subunits are incorporated into the complex. We hypothesized that assembly would stall at the step when the downregulated subunit would be inserted into the complex. We decided to immunocapture putative intermediates by C-terminal HA-tagging of one of the two core proteins, UQCRC2 (Fig. 5A). We assumed that the *T. brucei* core proteins would heterotetramerize into a module early during Complex III assembly, as in yeast (Stephan & Ott, 2020), and thus be appropriate for capturing putative sub-assemblies.

**Figure 5.**
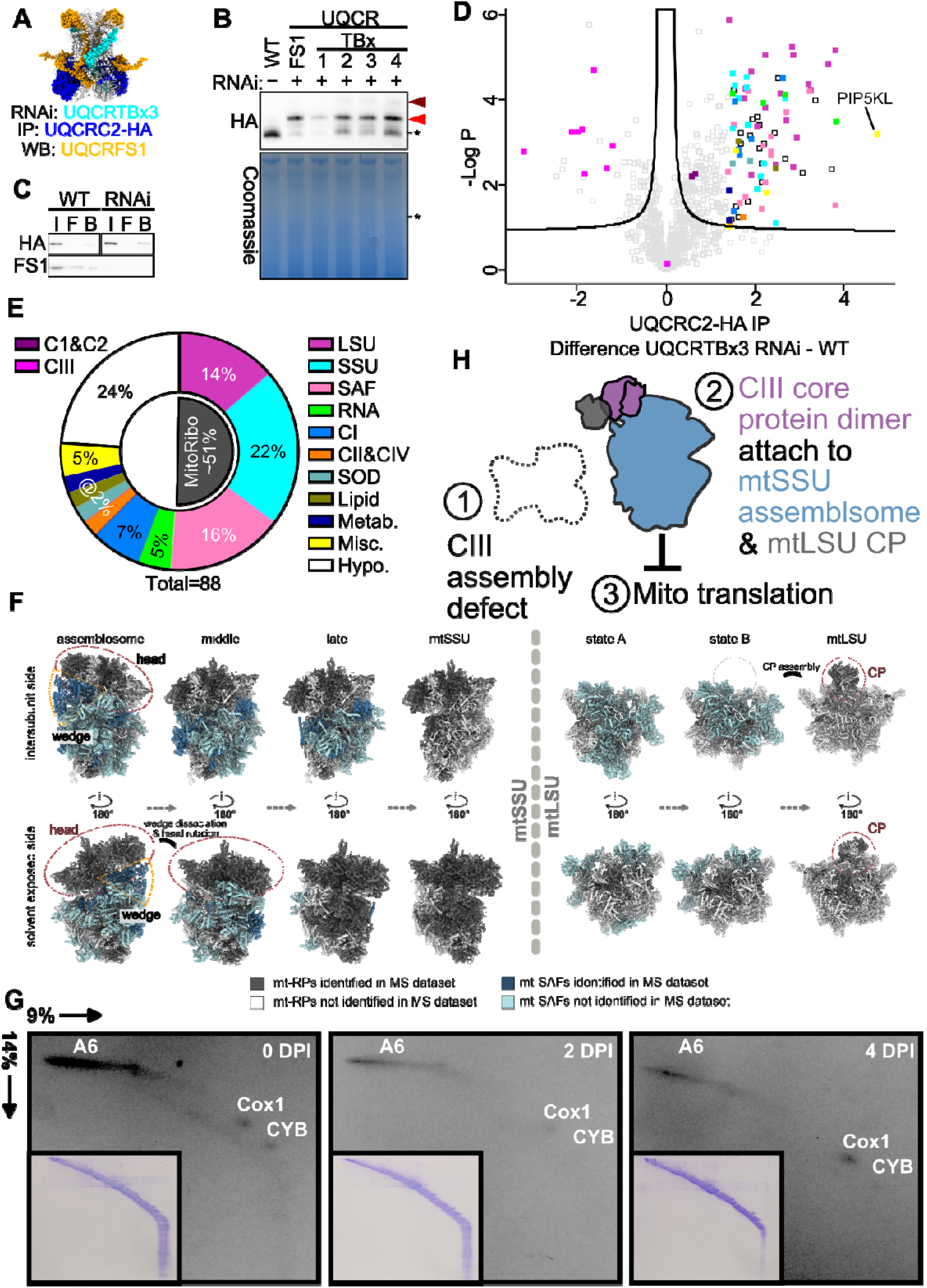
Ubiquinol: cytochrome c oxidoreductase assembly impairment results in aggregation of UQCRC2 and mitoribosome intermediates. **(A)** Key Complex III subunits for experimental design mapped onto structure from Fig. 3D. UQCRTBx3 (RNAi target) in turquoise, UQCRC2 (immunoprecipitation bait) in dark blue and Rieske protein (UQCRFS1) in yellow. (B) BN PAGE and western blot detecting UQCRC2-HA in UQCRTBx1-4 (Tbx1-4) or UQCRFS1 (FS1) RNAi background. Cell lines were induced for four days. The Coomassie-stained gel is shown for loading. Dark and light red arrowheads point to faint larger and strong smaller >720 kDa (*) bands that appear during RNAi-silencing (+) of aforementioned subunits. (C) Western blots showing successful immunoprecipitation of UQCRC2-HA with antibody recognizing epitope (HA) and Rieske (FS1) in input (I), flow thru (F) and HA-capturing bead (B) fractions derived from wild-type (WT) and UQCRTBx3-depleted (RNAi) cells **(D)** Volcano plot showing enriched proteins in UQCRC2-HA immunoprecipitation (IP) in UQCRTBx3 RNAi versus WT samples color-coded as in E. Note that Complex III subunits are depleted in this experiment due to UQCRTBx3 RNAi (see Figure 3). PIP5KL, phosphatidylinositol-4-phosphate 5-kinase-like; x-axis, Log_2_-transformed fold-enrichment; y-axis, -Log-transformed P-values from triplicate measurementI**(E)** Pie chart depicting the composition of >1.4-fold enriched proteins in D. Color-coding key (shared with D) to right. MitoRibo, mitoribosome; LSU, large subunit; SSU, small subunit; SAF, SSU assembly factor; RNA, RNA metabolism proteins; CI, Complex I; CII, Complex II; CIV, Complex IV; SOD, Superoxide dismutase; Lipid, lipid metabolism protein; Metab., metabolism enzyme; Misc., miscellaneous; Hypo., hypothetical protein. See table S3. Top left, color coding only in D. C1 & C2, UQCRC1 and UQCRC2; CIII, Complex III. **(F)** 1.4-fold enriched mitoribosomal proteins (mt-RPs) mapped onto solved structures (adapted from (Gahura *et al*., 2022a). CP, central protuberance. **(G)** Exposure of two-dimensional gels to visualize metabolically ^35^S-labelled *de novo* synthesized mitochondrial polypeptides in WT (DPI, 2 and 4-day post-induction (DPI) of RNAi of UQCRTBx3, labelled 0, 2, AND 4 DPI, RESPECTIVELY. Spots corresponding to ATP synthase subunit a (A6), Cytochrome c oxidase subunit 1 (Cox1), Cytochrome b (CYB) are labelled in each image. Insets are Coomassie stained two-dimensional gels as a loading control. Note background labelling by residual cytoplasmic translation after cycloheximide inhibition overlays the Coomassie-stained. *, unincorporated ^35^S methionine. Representative images from biological triplicates were exposed on the same film. **(H)** Hypothetical model of mitoribosome aggregation with the UQCRC1/2 heterodimer upon impairment of Complex III assembly.

UQCRC2-HA was introduced into each existing cell line for inducible RNAi of 4 UQCRTBx subunits or UQCRFS1. However, we did not observe various subassemblies corresponding to stalling at each downregulated subunit’s insertion as originally hypothesized. Instead, we observed that most UQCRC2-HA incorporated into the same two >720 kDa assemblies, which are larger than Complex III, when any of the assayed subunits were silenced (Fig. 5B).

This finding prompted us to identify the proteins that associate with UQCRC2-HA when Complex III biogenesis is disrupted. We decided to immunoprecipitate UQCRC2-HA in wild-type and UQCRTBx3 RNAi cell lines. As expected, UQCRFS1 was not detected after RNAi, but was immunocaptured by UQCRC2-HA under wild type conditions (Fig. 5C). The beads from both immunoprecipitations were analyzed by mass spectrometry and 88 proteins were identified to associate with UQCRC2-HA during UQCRTBx3 RNAi (Fig. 5D). Conspicuously, more than half of this cohort are proteins related to mitochondrial ribosomes (Fig. 5F).

Specifically, we identified 19 and 12 mitoribosomal proteins (mtRPs) of mtSSU and mtLSU, respectively (Ramrath *et al*, 2018). We also detected 14 assembly factors (AFs), which have been previously found in at least one precursor of the small subunit (Gahura *et al*, 2022a; Lenarčič *et al*, 2022; Saurer *et al*, 2019; Zgadzay *et al*, 2025). Spatially, 11 out of 19 mtRPs are present in the head of the mtSSU. A cluster of five mtRPs localize onto the solvent exposed surface of mtSSU and are recruited relatively late in the assembly process (Lenarčič *et al*., 2022). All but one of the identified AFs are present in the earliest characterized mtSSU precursor termed the assemblosome (Saurer *et al*., 2019; Zgadzay *et al*., 2025). Seven of these constitute most of a distinct structure of nine AFs wedged between the head and body, keeping a compositionally mature head in a rotated immature position (Saurer *et al*., 2019). Five out of 12 pulled-down mtLSU mtRPs are recruited very late during mtLSU biogenesis (Fig. 5E) to constitute almost the whole central protuberance (CP) (Jaskolowski *et al*, 2020; Tobiasson *et al*, 2021). The CP is a functionally conserved structural element of all ribosomes that forms multiple inter-subunit bridges with the mtLSU head during translation.

Because dissociation of the wedged cluster of AFs is key for subsequent steps in mtSSU biogenesis, we surmise that the immunocaptured core protein/mtSSU-precursor admixture may serve as a Complex III assembly checkpoint. Furthermore, maturation of CP might provide yet another means of interplay between Complex III and mitoribosome assembly. These possibilities led us to hypothesize that mitochondrial translation is affected upon UQCRTBx3 downregulation. To test the null hypothesis, we metabolically labelled *de novo* translated mitochondrial polypeptides (Hashimi *et al*, 2013; Skodova *et al*, 2013) in WT and UQCRTBx3 knockdowns 2 and 4 days after induction (Fig. 5G). All translation products whose identity is known were decreased upon UQCRTBx3 depletion. The signal corresponding to MT-CYB, the mitochondria-encoded subunit of Complex III, was particularly diminished. Thus, we conclude that mitochondrial translation is suppressed when assembly of Complex III is impaired.

Based on our results, we propose that failure to assemble nucleus-encoded subunits may result in trapping of the assemblosome onto the core proteins to downregulate futile mitochondrial translation (Fig. 5G). Alternatively or additionally, the translating mitoribosome may crosstalk with Complex III assembly *via* contacts in the region containing the mtSSU head and mtLSU CP. The role of unusually abundant *T. brucei* mitoribosomal assembly intermediates as checkpoints has been proposed previously (Gahura *et al*., 2022a; Lenarčič *et al*., 2022), and biogenesis of mitoribosomes in general is responsive to the physiological state of mitochondria (Brischigliaro *et al*, 2024). Translation regulation of specific mitochondrial transcripts in response to a supply of nuclear encoded subunits of respective OPXHOS complexes has been documented in model eukaryotes (Carlström *et al*, 2026; Herrmann *et al*, 2013; Moran *et al*, 2024; Richter-Dennerlein *et al*, 2016; Soto *et al*, 2022). Our data here leads us to hypothesize that OXHPHOS biogenesis can also provide feedback to finetune mitoribosome biogenesis to mitigate an imbalance of mitochondria- and nucleus-encoded subunits of Complex III. In the context of our study, depletion of any of the nucleus-encoded Complex III subunits may have shifted this putative finetuning mechanism toward accumulation of mitoribosome intermediates on the UQCRCP2-HA.

Finally, we point out that the most enriched co-IP protein in this experiment is a phosphatidylinositol-4-phosphate 5-kinase (PIP5K) like protein (Tb927.1.740) (Fig. 5D). This class of kinases are known to be involved in the biogenesis of lipid-derived secondary messengers that regulate vesicular trafficking, cell cycle progression and differentiation (Porciello *et al*, 2016). However, no PIP5K like protein has been reported to be imported into the mitochondrial matrix or directly regulate any aspect of mitochondrial gene expression. Thus, this observation brings another dimension into kinetoplastid–and perhaps other eukaryotic–mitochondrial biogenesis. I

In conclusion, our complexome data has led to the serendipitous discovery of a link between Complex III assembly and mitochondrial translation. We show that ablation of an integral subunit of Complex III leads to suppression of mitochondrial translation. This also demonstrates that exploration of our complexome data can lead to establishing exciting lines of future research.

### Two subunit g-like proteins associate with F_o_F_1_-ATP synthase monomers

During the assessment of Complex V subunit profiles, we noticed two proteins that appeared to interact almost exclusively with F_o_F_1_-ATPase monomers, which is highly unusual for the known subunits of the complex (Fig. 6A). These proteins were dubbed g-like’ (g’) and g-like’’ (g’’) due to the similarity of their primary and AF3-predicted tertiary structures with the *T. brucei* g-subunit (Fig 6B, Fig S6A, Table S4), a key subunit for Complex V dimerization (Gahura *et al*., 2022b). The kinetoplastid subunit g and g-like proteins likely share a common origin as they form a well-supported monophyletic group in a maximum likelihood phylogenetic tree (Fig. S6B).

**Figure 6.**
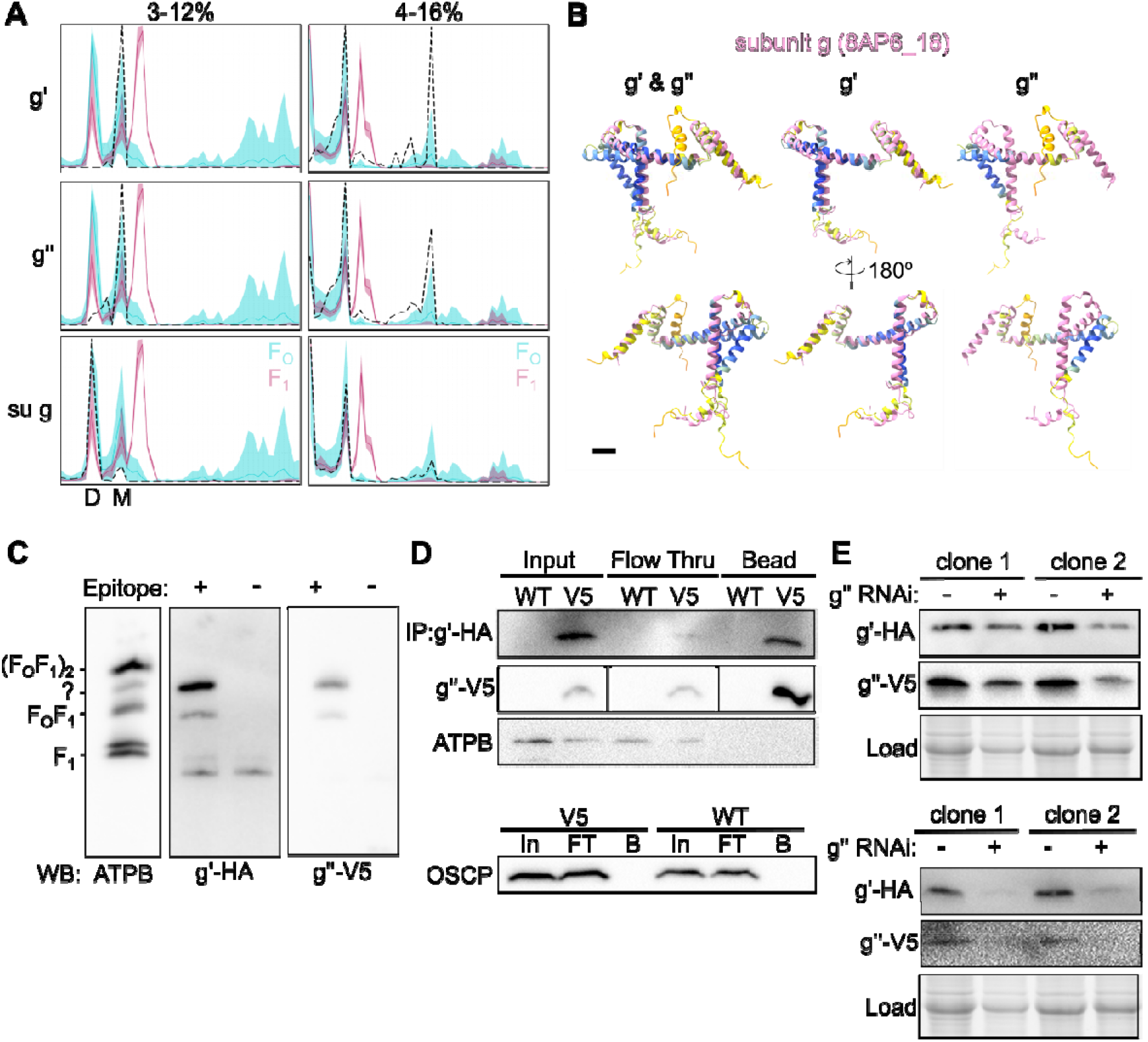
Subunit g paralogs form a heterodimer and are not assembled into F_O_F_1_-ATP synthase dimers. **(A)** The relative abundancies of g-like proteins g’ and g’’, plus subunit (su) g (dashed black line) across the 48 size fractions in PCF. As in Figure 1B. Color coding key on top right of su g plots. Dimer ((F_O_F_1_)_2_) and monomer (F_O_F_1_) peaks indicated by ‘2’ and ‘1’, respectively, at bottom of su g plots. **(B)** Structural alignment of AlphaFold 3 predicted g’ and g’’ (using standard local pLDDT confidence score coloring (Rennie & Oliver, 2025)) with solved su g (in pink) from (Gahura *et al*., 2022b). **(C)** Western blot (WB) of BN PAGE resolved cells without (-) or with (+) epitope tagged g-like proteins probed with antibodies recognizing F_1_ ATP synthase subunit β (ATPB), band possibly corresponding to CIICIV_2_CV supercomplex that incorporates the g-like proteins. **(D)** Immunoprecipitation (IP) of g’’-V5 co-IPs g’ HA but not ATPB or oligomycin-sensitivity conferring protein (OSCP) subunits of F_0_F_1_-ATP synthase. WT, wild-type negative control; In, input; FT, flow thru; B, beads. **(E)** Two clones of g’ and g’’ inducible RNAi (+) cell lines probed with antibodies recognizing epitopes on left. Part of the stain-free gel is shown as a loading control.

To further investigate, we epitope tagged g’ and g’’ with the HA and V5 epitopes, respectively. We leveraged these tags in BN-PAGE to confirm if they migrate with the F_o_F_1_-ATPase monomers, as was observed in the complexome profiles (Fig. 6A). A minor portion of g-like proteins co-migrated with the F_o_F_1_-ATPase monomers on BN-PAGE (Fig. 6C). But unexpectedly, most of the g’ and g’’ associated with a weak ATPB immunosignal between the F_o_F_1_-ATPase dimer and monomer. This implies that the band represents a different entity than the F_o_F_1_-ATPase monomers. We speculate that the reason this band was not detected in our complexome datasets is because the BN PAGE used a different detergent to solubilize the native complexes prior to electrophoresis. The former employed dodecyl maltoside (DDM) whereas the detergent for the latter was digitonin.

We also exploited these tags to verify that ǵ and g’’ interact with each other (Fig. 6D). However, under our IP conditions, we failed to detect co-IP of the ATPB, one of the two catalytic core subunits in the F_1_ head of Complex V, or the F_o_ subunit Oligomycin Sensitivity-Conferring Protein (OSCP). This implies that the g-like proteins heterodimerize but do not interact strongly with these Complex V subunits.

To facilitate functional analysis, this cell line mentioned above was transformed with RNAi constructs downregulating individual or both g-like proteins (Fig. S6C). Induction of RNAi did not affect the growth of any of the cell lines in media that promotes respiratory growth (Fig. S6D), perhaps owing to the incomplete depletion of the targeted proteins (Fig. S6C). Furthermore, the assembly and levels of the major Complex V forms were not conspicuously affected by depletion of the g-like proteins (Fig. S6E). However, we did observe the stability of the g-like proteins are mutually dependent on each other, consistent with the robust interaction we observed in the IP experiment (Fig. 6E).

Using the complexome, we have found two g-like proteins that associate with F_o_F_1_-ATPase monomers as well as a slightly larger assembly of unknown origin that spuriously appears in our BN PAGE (*c.f.* Fig. 6C and Fig. S6E). During the course of this project, we have entered a collaboration further characterizing these g-like proteins. The major finding of this study is that the g-like proteins are required for the formation of a supercomplex made up of Complexes II, IV dimer and a F_o_F_1_-ATPase monomer, *i.e.* a CIICIV_2_CV (please consult Hu et al.(2026) for a more in-depth examination of this complex). We speculate that the unknown band appearing in Fig. 6C may represent this supercomplex, which is slightly larger than the F_o_F_1_-ATPase monomer. Here, we demonstrate that the g-like proteins mare stable only when they interact with each other. The intrinsic instability of monomeric g-like proteins may add an extra layer of regulation, protecting against spurious interaction with the canonical g subunit, a condition that could iinterfere with F_o_F_1_-ATPase dimer formation.

### A comparison of trypanosomatid complexomes

During the revision of this article, another complexome study analyzing the same two *T. brucei* life cycle stages we did plus *Leishmania tarentolae*, a trypanosomatid isolated from a gecko (Klatt *et al*, 2019), was published (Solayman *et al*., 2026). A logical question is how do our complexome data compare with theirs? Two key methodological considerations distinguish our complexosomes from theirs; that of the number of slices per gel assessed (48 vs 60) and importantly, the detergent employed to lyse the cells (digitonin vs DDM). All complexome datasets from both studies detected between 2000 and 3500 proteins. (Fig. S7A-B). There is an overlap among in the proteins detected in both studies. In total ∼1345 and ∼1130 PCF and BSF proteins, respectively, were detected in both datasets (Fig. S7C). Overall, we detected two thirds of proteins identified in the Solayman complexomes. Conversely, they detected just under half of proteins found in our study.

Both studies captured a similar number of mitochondrial proteins (Fig. S7A). However, there is less agreement between the mitoproteomes captured in the two studies (Fig. S7D) than the detected total protein (Fig. S7C). However, this is advantageous as the union of detected mitochondrial proteins from both studies has higher coverage of the 1666 proteins that comprise the mitoproteome (Vacas *et al*, 2026) than each complexome alone (Fig. S7D). It should be noted that the number of shared and unshared proteins is likely to be under and overestimated, respectively, in our meta-analyses. This is in large part due to differences in the system used to assign proteins to groups of paralogs (see Material and Methods for an in-depth explanation).

Some key technical differences potentially underlie other disparities between the two studies (Table 2), The different organelle enrichment protocols may explain why complexome data from Solayman et al. show better coverage of the mitochondrial proteome (mitoproteome) (Vacas *et al*., 2026) than ours (Fig. S7E), reflecting the higher purity of mitochondria derived from their isolation protocol. Conversely, our complexomes captured a higher portion of the glycosomal proteome (Güther *et al*, 2014) (Fig. S7F), to a similar degree as a previous study (Peikert *et al*, 2017). This suggests that the lesser representation of the mitoproteome in our datasets is balanced by a greater share of non-mitochondrial complexes.

**Table 2.** Technical differences in how complexome prepared between our study (Benz) and that of Solyman (2026).

| Technical detail | Benz | Solayman |
|---|---|---|
| Enrichment of biomass to produce sample for Complexome analysis | **Organelles.** Permeabilization of cells with 0.15 % (w/v) digitonin yields indiscriminate enrichment of organelles, leading to a wider but shallower sampling of their respective proteomes<sup>1,2</sup>. | **Mitochondria.** Density gradient sedimentation to more stringently separate mitochondria from other organelles, leading to deeper and more precise sampling of the mitoproteome<sup>2</sup>. |
| Detergent for biomass solubilization prior to BN-PAGE | 1.5% (w/v) digitonin. | 2:1 n-dodecyl-β-D-maltoside (DDM):protein. |
| Number of fractions (*i.e.* number gel slices after BN-PAGE | 48 | 60 |

<sup>1</sup>(Peikert *et al.*, 2017)
<sup>2</sup> (Niemann *et al*, 2013)

This difference in organelle enrichment may also explain why in our study, there is considerably less of the mitoproteome detected in BSF versus PCF (Fig. S7G-H). In BSF, mitochondria occupy an smaller portion of the cell body than in PCF (Bílý *et al*., 2021; Zíková, 2022). This would be reflected in less mitochondria represented in the crude organellar enrichment we performed. Nevertheless, Solayman and colleagues also detect less of the mitoproteome in BSF than PCF (Solayman *et al*., 2026; Vacas *et al*., 2026) (Fig. S7G-H). This suggests perhaps a reduced protein complex landscape in BSF mitochondria despite the smaller than expected difference in mitoproteome complexity (Fig. S7H) (Vacas *et al*., 2026).

We benchmarked the compatibility of our complexome data with Solayman and colleagues’ by investigating whether the singular MCP we discovered to interact with Complex I beta (Tb927.6.1630, Fig. 2) co-fractionated in their datasets. Indeed, these molecules exhibited similar profiles (Fig. 7I). However, these profiles were considerably more dispersed than the profiles we observed (Fig. 2A). This may be due to either the different detergents and/or number of fractions employed in the respective studies (Table 2). The ortholog of this MCP (LtaP30.0290) was not detected in the *L. tarentolae* complexome. Furthermore, Complex I subunits and the MCPs exhibited a fractional dispersal that precludes detection of co-fractionation among these molecules (Fig. 7I). Thus, whether this interaction we found in *T. brucei* is conserved among trypanosomatids remains unknown at present.

We ran GIP analysis on the three Solayman complexomes (Table 1, S1) to further test the compatibility of both studies. Specifically, we compared the number of proteins that were unbiasedly assigned to a GIP cluster, which relies on the occurrence of proteins with unequivocally correlated distribution profiles. The fraction of proteins placed in GIPs in our study ranged from 12 to 20.6%, which is a less variable distribution than was produced with the Solayman data (Fig. S8A). The outlier that is responsible for the wider standard deviation is the from Solayman BSF data, in which only 8% of detected proteins were assigned to a GIP group. This is less frequent than from any of our complexomes (Fig. S8A), which is unexpected given that an equivalent number of proteins were detected in all complexomes from both studies (Fig. S7B). Otherwise, the largest portion of proteins were assigned to GIP groups from the rest of the Solayman datasets, which also exhibited a narrow distribution (Fig. S8A). The same general trend was seen for in the amount of mitochondrial proteins that were sorted into a GIP group (Fig. S8B), with two exceptions: (1) A more sporadic distribution of GIP assignment rates than for total protein (Fig. S8A); (2) the outlier comes from the extraordinarily lower rate of GIP assignment of 9.8%. The latter could be due to the more dispersed profiles observed in the *L. tarentolae* (S7I), which would confound GIP grouping.

A minority of proteins placed into GIP groups were shared in the two studies (Fig. S8C) Roughly one third of our and a quarter of their GIP proteins intersected in PCF. Our respective GIP protein sets overlapped to a lesser degree in BSF. This could stem from many factors, such as the high number of proteins that were only detected in one study (S7C). Other technical details could underlie the disparity in proteins placed in GIP groups in both studies, especially the different number of fractions analyzed (Table 2).

To address the overlap of proteins incorporated into GIP clusters in both studies, we compared the predictions made by each GIP grouping to the polypeptide composition of the solved structure obtained from *L. tarentolae* (Fig. S8D, Table S1) (Solayman *et al*., 2026). All *T. brucei* GIP groups accurately predicted Complex III subunits. However, virtually all of these GIP groups contained one false positive. UQCRH is omitted in the GIP predictions from our PCF 4-16% and union datasets. The *L. tarentolae* GIP-grouping also correctly recapitulates Complex III. However, an additional ten superfluous false positives were also placed into this GIP group. In conclusion, GIP-groupings made from complexome datasets from both studies were mostly in agreement regarding Complex III, with a high confidence reconstitution can be derived from the intersection of all these datasets (*e.g.* Fig. 3D).

Interestingly, the UQCRC1/2 heterodimer is rendered as the independent GIP group 1174 (Fig. S8D). This GIP-grouping is consistent with the apparent functional separation of the heterodimer from the other subunits (*e.g.* Fig. 5), likely stemming the import of the former’s import into the matrix, in contrast to the other subunits (Mach *et al*, 2013). Conditions used in the 4-16% BN PAGE experiment seem to have preserved the semiautonomous nature of the UQCRC module This may have led to a moderate deviation of the heterodimer’s fractional distribution profile compared to the other Complex III subunits (Fig. S8E), which was detected by the GIP algorithm.

We conclude from these analyses that the complexomes presented by us here and Solayman and coworkers are complementary. The unbiased GIP groups can be compared across datasets, allowing hypotheses to be addressed *in silico* prior to lab work. Furthermore, the stability of protein complexes in the different detergents used in both studies can be leveraged, bringing added value to these datasets.

## Conclusions

### Summary of findings

Here, we have implemented complexome profiling in the model euglenozoan *T. brucei* to observe the mitochondrial multiprotein complex landscape of two life cycle stages. Collectively, these data along with those derived from the apicomplexans plasmodium (Evers *et al*., 2021) and toxoplasma (Maclean *et al*., 2021), plus an oyster parasite related to the the former two species(Maclean *et al*, 2026), bring needed diversity to the complexome profiling ecosystem.

Roughly 85% and 55% of the estimated 1120 proteins imported into the mitochondrion (Peikert *et al*., 2017) were detected in our PCF and BSF complexome profiles, respectively. We have analyzed this data to find a MCP protein associating with a Complex I module, a heterodimer of subunit-g derived proteins that are at the interaction hub of an unprecedented respiratory chain supercomplex, a likely Complex III assembly factor, and uncovered a link between Complex III assembly with mitochondrial translation. Taken together, our findings demonstrate the utility of complexome datasets in generating hypotheses that have revealed multifarious interconnections that exist among the respiratory chain complexes, including expression of their mitochondria-encoded subunits.

Our dataset is not restricted to mitochondrial complexes: 48-65% of proteins expressed in each life cycle stage (Moloney *et al*, 2023) are present in our complexome profiles. Thanks to our organelle enrichment protocol, we have captured 47-61% of the 129 proteins that make up the high confidence glycosomal proteome (Güther *et al*., 2014), similarly to previous studies (Peikert *et al*., 2017). Thus, we hope that our complexome profiling datasets will be a suitable addition to the plethora of useful resources generated by the dedicated researchers in the kinetoplastid field.

### How to use the complexome data in your research

The unbiasedly assigned GIP groups of proteins with correlated profiles are available for perusal (Table S1). All individual and combined BSF and PCF GIPs are listed in parallel to facilitate comparison. Furthermore, KEGG annotations (Kanehisa *et al*, 2016) of each protein are provided for end user annotation. Unfortunately, the Nova software that clusters and visualizes proteins based on their profiles (Giese *et al*, 2015) is currently unavailable.

However, we have overcome this by uploading our complexome datasets (Dataset S1) to the spatial proteomics fraction profiling tool pRoLoc (Crook *et al*, 2019) to generate tSNE and distribution plots, such as those depicted in Figs. 1A and B, respectively. In validating any groupings, one should consider that false negatives, *i.e.* the failure to detect interacting proteins by complexome profiling, are almost certainly present in our dataset, as is in any large dataset. However, false positive interactions are easy to test using the streamlined tagging developed in *T. brucei* (Billington *et al*, 2023; Dean *et al*, 2015) we implemented here to verify the interactions implied in the complexome profiling data.

Our complexomes are complementary to those reported by Solayman and colleagues (2026). Used together, they can be used to explore the protein complex landscape in *T. brucei*. While we predominantly use combined complexome datasets for streamlined analysis in this study, there are certain protein interactions that can be lost if the individual plots are also not consulted in tandem. Thus, we encourage researchers to examine the behaviour of their proteins-of-interest across both individual and combined datasets, for a comprehensive understanding of protein complexes in *T*. *brucei*.

## Materials and Methods

### Culturing and transformation of cell lines

The *Trypanosoma brucei* strains SmOxP9 and SmOxB4, derivatives of PCF 927 and BSF427 strains (Poon *et al*, 2012), were used in this study SmOxP9 tutively expresses the tet repressor and T7 polymerase, thus was used for all genetic manipulations. Cells were regularly grown in SDM79 but shifted to glucose-lacking SDM80 (Coustou *et al*, 2008) for growth analysis. Antibiotics used were 1 μg/ml puromycin (for the parental strain SmOxP9), 50 μg/ml hygromycin (for V5 tags), 15 μg/ml G418 (for HA tags) and 2.5 μg/ml phleomycin (for RNAi cell lines).

### Generation of PCR products for tagging and plasmids for RNAi

Epitope tagging was performed as described in (Dean *et al*., 2015) using long primer PCRs to append V5 or HA tags (Kaurov *et al*., 2018) to the C-terminus of the proteins of interest. 90% of a 50 µl PCR reaction performed with 20 ng of template plasmid was directly used for transfection without prior purification (see below). For RNAi-mediated depletion, fragments of TBx1-4 (Tb927.3.700, Tb927.11.16510, Tb927.10.4240 and Tb927.5.2560), Rieske protein (Tb927.9.14160), g’ (Tb927.11.10040) and g’’ (Tb927.7.7440) designed by RNAit (Redmond *et al*, 2003) into the expression vector p2T7-177 (Wickstead *et al*., 2002), using BamHI and XhoI restriction sites. To generate the vector targeting both g’ and g’’ simultaneously, a fragment of g’’ was cut out of the existing p2T7-177 + g’’ vector using an internal BglII site in combination with BamHI. The p2T7-177 + g’ RNAi vector was 16inearized with BamHI, dephosphorylated and subsequently ligated to the BamHI-BglII fragment of g’’. All RNAi constructs (about 12 µg of plasmid) were 16inearized with NotI prior to transfection. Primer sequences are listed in Table S5.

### Transfection of T. brucei

5x10^7^ cells from a logarithmically growing culture were pelleted, resuspended in 100 μl of transfection buffer (Schumann Burkard *et al*, 2011) and mixed with either PCR products (for epitope tagging) or linearized plasmids (for RNAi). An Amaxa Nucleofector 2b (Lonza) with program X-014 was used for electroporations and the cells transferred to SDM79 without antibiotics for recovery overnight. Selective antibiotics were added the following day (concentrations see above) and the cells left to recover for 10-14 days.

### Growth curves

For growth analysis, the respective RNAi cultures were grown in triplicate in SDM80 medium in the presence or absence of 1 µg/ml doxycycline, a compound that induces RNAi (Wickstead *et al*., 2002). Cell densities were determined using the Beckman Coulter Z2 Cell and Particle Counter every 24 hours and cultures diluted back to the starting density of 1 × 10 cells/ml every 48 h to maintain exponential growth.

### Sample preparation and blue-native polyacrylamide gel electrophoresis for subsequent complexome analysis

PCF and BSF *T. brucei* were permeabilized with 0.015% (w/v) digitonin and subsequently centrifuged to separate the mitochondria-enriched organellar fraction from the cytosol as performed previously (Sheikh *et al*., 2025). The organellar pellet was solubilized in 1.5% (w/v) digitonin in dissolved in 1X NativePAGE (Invitrogen) sample buffer and 0.05 U Dnase I (ThermoFischer) and incubated for 1 hour on ice. The lysate was clarified by centrifugation at 21,000 × g for 45 min at 4 °C. The protein concentrations of the lysates were measured using a Bradford assay, and 50 μg total protein was combined with 5% Coomassie brilliant blue G-250 prior to loading onto a 3–12% or 4-16% Bis-Tris NuPAGE™ Bis-Tris Mini Protein gel (Invitrogen). Electrophoresis was carried out for 2.5 h at 150 V at 4 °C, after which the gel was washed twice in ddH₂O for 5 min each. The gel was then destained in a solution containing 50% methanol and 10% acetic acid until excess Coomassie dye was removed, followed by two additional 5 min washes in ddH₂O.

### Complexome sample preparation and mass spectrometric analysis

Each lane was excised from the BN PAGE gel and subsequently cut into 48 1.25 mm slices. Slices were cut into ∼1 mm^3^ cubes and transferred to a 96-well filter plate (Multiscreen Solvinert MSRLN0450, 0.45 µm pore size PTFE membrane, Merck Millipore). In-gel digestion was performed according to a protocol (Shevchenko *et al*, 2006) modified for a 96-well filter plate processed on a vacuum manifold. Briefly, reduction by DTT was followed by iodoacetamide alkylation, de-staining, and trypsin digestion. Tryptic peptides were extracted, dried, dissolved, desalted by 1-layer C18 StageTips, and transferred to a U-shaped 96-well plate suitable for the Ultimate 3000 nano UHPLC autosampler.

Peptides were separated on a 15 cm C18 column using 30 min elution gradient and analyzed in a DDA mode on Orbitrap Exploris 480 MS equipped with a FAIMS unit set at CV of -45 V. Raw files were processed in MaxQuant (v. 2.7.3.0) using following protein fasta files: Trypanosoma brucei proteome TriTrypDB-68_TbruceiTREU927_AnnotatedProteins.fasta downloaded from https://tritrypdb.org and custom fasta Tbrucei_mtEncoded_OG_2024-06.fasta containing mitochondrial DNA-encoded proteins compiled from https://kdna.net/trypanosome/seqs/index.html. Resulting iBAQ values (proteinGroups.txt report file) of each protein detected in the individual slices were normalized to the protein with the highest intensity. The resulting data files for BN PAGE-resolved proteins from PCF and BSF are referred to as PCF107 and BSF107 as well as PCF139 and BSF139 for 4-16% and 3-12% BN PAGE gels, respectively.

### Unbiased protein complex identification using Gaussian Interaction Profiler

Protein complexes were identified by the following strategy. The ProteinGroups.txt was first filtered to remove protein IDs which were contaminants, identified by site or reverse hits only. Following this, only the ID names and IBAQ values per slice were retained. The IBAQ values were normalized to the relative value of 1 per sample, and the following samples were combined – PCF107 and BSF107 were combined into a single sheet, and PCF139 and BSF139 were combined into a separate single sheet. The 2 final datasets were run through the Gaussian Interaction Profiler (GIP) (van Strien *et al*., 2024), via the GIPit colab implementation written for this study and deposited in GitHub (https://github.com/Rayyan-Tariq-Khan/GIPit?tab=readme-ov-file). The following parameters were used: cluster ratio 0.5 and 4 bootstraps. The resulting output was then used to determine the complexing behavior of the proteins.

### Homology searches and AlphaFold3 modeling

The UBQCR complex sequences listed in Table S1 were entered into the AlphaFold3 server. Two copies of each subunit were modelled except UBQRB due to limitations in tokens (*i.e.* amino acid residues) that can be handled for one multi-protein structure. The resulting Crystallographic Information Files (CIFs) were visualized in ChimeraX 1.10 (Meng *et al*, 2023). For structural alignment of UQCRTBx subunits to putative *E. gracilis* orthologs, the Matchmaker command performed in default setting using a given UQCRTBx to query the whole solved *E. gracilis* Complex III complex (PDB 8IUF). This ChimeraX tool was used to align AF3-predicted g’ and g’’ structures to subunits from the solved *T. brucei* Complex V structure (PDB 8AP6). AlphaBridge software (Álvarez-Salmoral *et al*, 2024) was used to map each binary interaction among the subunits in the predicted Complex III structure and local pLDDT scores mapped onto the primary structure of each predicted subunit. Other structural homology searches were performed using FoldSeek (van Kempen *et al*., 2024). The predicted AlphaFold structure of *T. brucei* LYRM7 was retrieved from a dedicated database of predicted kinetoplastid protein structures *(Wheeler, 2021)*.

Kinetoplastid subunit g and g-like sequences listed in Table S4 were retrieved from the TriTrypDB database (Aslett *et al*, 2010). Sequence alignment was performed using the MAFFT algorithm (Katoh *et al*, 2002) implemented in Jalview (Waterhouse *et al*, 2009). Molecular phylogenetic analysis was performed using PhyML (Guindon *et al*, 2010) with LG model of amino acid substitution and 100 bootstraps. As an outgroup, putative subunit g sequences from diplonemids, the sister group to kinetoplastids (Kostygov *et al*., 2021), was retrieved by BLASTP using kinetoplastid subunit g sequences as a query in EukProt v3 (Richter *et al*, 2022).

### Immunoprecipitation of UQCRCP2-HA and identification of co-purifying proteins by mass spectrometry

TBx3 RNAi cell lines were either induced or not for 4 days preceding immunoprecipitation. About 2 x 10^8^ cells were separated into cytoplasmic and organellar fractions as previously described using 0.015% digitonin (Kaurov *et al*., 2018). The pellet containing the organellar fraction was subsequently lysed in mitochondrial lysis buffer (20 mM Tris-HCl (pH 7.4), 50 mM NaCl, 10% (v/v) glycerol, 0.1 mM EDTA, 1 mM PMSF, 1.5% digitonin) on ice for 1 h. Following centrifugation at 16,000 × g for 30 min at 4°C, the supernatant was collected as the cleared lysate. An aliquot was retained as the input (IN) fraction, and the remaining lysate was incubated with HA-coupled paramagnetic beads (ThermoFisher) for 3 h at 4°C with rotation. The unbound material was then collected as the flow-through (FT) fraction and beads were washed five times with lysis buffer (the last two washes omitting the detergent). An aliquot of the beads was retained as the eluate (EL) for analysis by SDS-PAGE and the remaining beads were stored at -80°C until mass spectrometry analysis. All IPs were performed in triplicate and every replicate analyzed by western blotting to confirm successful capture of the tagged bait.

#### SDS PAGE and western blotting

Protein samples were separated using home-made 12% SDS-PAGE gels containing 0.5% (v/v) 2,2,2-trichloroethanol allowing for total protein visualization via UV exposure. The proteins were then transferred onto PVDF membranes (Amersham), and probed with the appropriate primary antibodies diluted in 5% milk (w/v) in PBS: HA (1:1,000, Sigma), V5 (1:1,000, Invitrogen), Rieske (UQCRFS1; 1:1,000) and TrCOIV (1:500) (Doleželová *et al*, 2020), plus ATP synthase subunit β, also known as ATPB (1:1,000) (Gahura *et al*, 2018). Secondary HRP-conjugated anti-rabbit or anti-mouse antibodies (1:2,000, Sigma) were used. Proteins were visualized using the Pierce ECL system on a VILBER Fusion instrument (VILBER). Precision Plus Protein All Blue Standard (Bio-Rad) was used as molecular weight marker. Blots shown are representative of at least 2 independent experiments.

#### BN PAGE analysis

Mitochondria-enriched fractions were solubilized in 1 × NativePAGE™ Sample Buffer (Thermo Fisher) containing 2% n-Dodecyl-β-D-maltoside (Sigma), 0.1 mg/ml DnaseI, cOmplete™ Mini Protease Inhibitor Cocktail (Roche) and 1 mM PMSF (Sigma) for 1 h on ice. Following centrifugation at 21,000 × g for 30 min at 4°C, the protein concentration was determined using a NanoDrop One (Thermo Fisher) by measuring absorbance at 280 nm. Approximately 70 µg of each sample was loaded onto a precast Native PAGE 3-12% Bis-Tris Gel (Invitrogen) and run for 2.5 h at 4°C, 150V. Proteins were blotted onto a PVDF membrane (Amersham) by wet transfer for 2 h at 4°C, 100 V. Following a blocking step with 5% (w/v) milk in PBS, the blots were incubated with primary antibodies: V5 (1:1,000, Sigma, produced in rabbit), HA (1:500, Invitrogen, produced in mouse), ATP synthase β (ATPB; 1:1,000) or Rieske (UQCRFS1; 1:200) at 4°C overnight. Secondary antibody incubation and visualization of signals was performed as described for SDS-PAGE. Blots shown are representative of at least 2 independent experiments.

#### Genomic DNA PCR

Genomic DNA from the parental SmOxP9 and the V5-tagged TBx2 cell lines was isolated using ExGene^TM^ Clinic SV kit (GeneAll) according to the manufacturer’s instructions. Primers were designed to anneal at the beginning of the open reading frame and 80 nucleotides downstream of the stop codon of TBx2, so that integration of the tagging cassette plus resistance marker could be easily distinguished by size from the PCR product obtained in the parental cell line. PCR was performed on about 20 ng of gDNA and the resulting products resolved on an 0.8% agarose gel.

### In vivo translation assay and two-dimensional electrophoresis

Products of mitochondrial translation were labeled in 2 x 10^7^ cells harvested from the culture in logarithmic phase of growth (5 minutes, 1000 x *g*, RT), washed twice in SoTEG medium (0.6 M sorbitol, 20 mM Tris-HCl, pH 7.5, 2 mM EDTA, 10 mM glucose), pelleted (5 minutes, 1100 x *g*, RT) and finally homogenized in 180 μl of SoTEG. Cycloheximide was added to the cell suspension to a final concentration of 200 μg/ml, respectively, to inhibit cytosolic translation. After 10 minutes of incubation at 27 °C with vigorous shaking, the products of mitochondrial translation were labeled *in vivo* for 1 hour with 20 μl of Met-^35^S-Label mix (Hartmann Analytic) per sample. Proteins were resolved in two-dimensional Tris-glycine-SDS PAGE (9% in the first dimension, 14% in the second dimension). Labeled products of mitochondrial translation were visualized by autoradiography as described previously (Škodová-Sveráková *et al*, 2015), and abundant proteins were visualized with Coomassie brilliant blue R250 staining.

#### Comparison of our and Solayman and colleagues complexome

To search whether individual proteins detected in our complexomes were also found in the Solayman et al. (Solayman *et al*., 2026) complexomes and recently established 1666 protein Mitocarta (Vacas *et al*., 2026), the list of accession numbers from our study (Dataset S1) were searched against the corresponding catalogs found in the supplementary information from the respective publications were compiled in Excel. The ‘COUNTIF’ function was used with each of our proteins as a query against the protein catalogs from the other studies. Protein lists from our and the Solayman et al., studies were used to query the catalog of high confidence glycosomes (Güther *et al*., 2014). The number of shared instances were counted. The limitation of this method is that the Solayman et al., protein catalog listed all paralogs in a single cell delimited by the character ‘|’ (*e.g.*’ Tb05.5K5.20|Tb927.5.4360’). Our dataset omits all but one of the paralogs (*e.g.* we only have ‘Tb927.5.4360’ in our datasets). Thus, these true matches would be false negatives using our strategy. This is responsible for the overestimate of unshared proteins in both sets being compared.

## Data availability

The mass spectrometry proteomics data have been deposited to the ProteomeXchange Consortium via the PRIDE partner repository with the dataset identifier PXD081276 (Complexome) and PXD082314 (UQCR2-HA IP).

## Author contributions

**CB,** Conceptualization, Formal Analysis, Investigation, Visualization, Writing – Original Draft Preparation; **RTK,** Data Curation, Formal Analysis, Methodology, Software, Writing – Review & Editing; **MH,** Formal Analysis, Visualization, Writing – Original Draft Preparation; **OG,** Formal Analysis, Funding Acquisition, Investigation, Visualization, Writing – Original Draft Preparation; **UK,** Investigation; LRC, Investigation, Writing – Review & Editing; **IŠ-S**, Investigation, Funding Acquisition; **PTB,** Software; **MV,** Conceptualization, Formal Analysis, Project Administration, Writing – Original Draft Preparation; **HH,** Conceptualization, Formal Analysis, Funding Acquisition, Visualization, Project Administration, Writing – Original Draft Preparation

## Conflict of interest statement

The authors declare no conflict of interest.

## Acknowledgments

We thank Alena Zíková (Biology Centre, Czech Academy of Sciences [BC CAS]) for antibodies, helpful discussions, and valuable advice, Carolina Hierro Yap (BC CAS) for expert guidance on BN PAGE, Atlanta Cook (University of Edinburgh) for guidance on structure software, and Julius Lukeš (BC CAS) for his support. This study was primarily supported by Czech Science Foundation (CSF) Grant 23-07674S to HH, and further by CSF grant 26-20429S and the project OP JAC CZ.02.01.01/00/22_008/0004575 RNA for therapy, co-funded by the EU, to OG, the project VEGA1/0709/26 to IŠS. We acknowledge the Proteomics Service Laboratory at the Institute of Physiology (supported by RVO, ID 67985823) and Institute of Molecular Genetics (supported by RVO, ID 68378050) of the Czech Academy of Science.

## Supplementary Figure Legends

**Figure S1. The migration of F_O_F_1_-ATP synthase during blue native polyacrylamide gel electrophoresis (BN PAGE) under conditions used for eventual complexome processing. (A)** Western blot (WB) using antibody recognizing F1 ATP synthase subunit β (ATPB) on left with image of Coomassie-stained gel on right. F_O_F_1_-ATP synthase speciesresolved on indicated at left of WB. **(B)** Size fraction distribution of F_O_F_1_-ATP synthase profiles color coded as indicated at top shown next to ATPB immunoblot from A.

**Figure S2. Complexome profiles of bloodstream and procyclic form *Trypanosoma brucei* from individual gels. (A)** The distribution of subunits of marker complexes (color-coded as in bottom key) of the complexome profiles from the individual BN-PAGE gels as indicated at the top of each t-SNE plot. **(B)** Selected Gaussian Interaction Profile (GIP) groups from combined complexome profiles plotted onto t-SNE plot in A. Group identification number of each group as listed in Table S1. See Figure 1 for t-SNE plots of complexome profiles from the combined BSF∪ and PSF∪ complexome profiles from each life cycle stage.

**Figure S3. Gaussian interaction profile (GIP) group metrics. (A)** The number of proteins assigned to a GIP group. Bar and whiskers, mean and SD, respectively. **(B)** The fraction peak of each GIP group. Bar and whiskers, median and IQR, respectively. **(C)** Size fraction calibration curve of PCF 3-12% BN PAGE gel using approximate molecular weights of F_O_F_1_-ATP synthase monomer (1048 MDa, CV), ubiquinol: cytochrome c oxidoreductase (720 kDa, CIII), succinate dehydrogenase (500 kDa, CII), and LYRM7 (22 kDa, LYM). Most GIP groups lie in area marked in grey.

**Figure S4. Additional data regarding UQCRTBx1-4 and LYRM7, four divergent components and putative assembly factor of ubiquinol: cytochrome c oxidoreductase. (A)** Venn diagram showing intersection of 3-12% PCF, 4-16% PCF, and PSF∪ GIP groups enclosing UQCRTBx1-4. (B) Growth of PCF when UQCRTBx1-4 is depleted by inducible RNAi (RNAi+) versus uninduced controls (RNAi-). Key in bottom right plot. Standard deviation at each time point (n=3) not visible at this scale. See D. (C) Demonstration by PCR that the V5-tagging cassette was integrated into only one allele of *uqcrtbx2.* Oligonucleotide annealing sites flanking the V5 cassette are red half-arrows above and below schema of *uqcrtbx2* locus. (D) Doubling time of cell lines depicted in B and F (grey). Bar and whiskers, mean and SD, respectively. (E) Multisequence alignment of pLYRM7 encoded in the genomes of the kinetoplastids listed on left. LYR motif and kinetoplastid (kin.) α-helix pair boxed. **(F)** Growth of PCF when LYRM7 is depleted by inducible RNAi (RNAi+) versus uninduced controls (RNAi-). Key on bottom right of plot. Standard deviation at each time point (n=3) not visible at this scale. See D. **(G)** Depletion of pLYRM7-V5 after a specified number of days post induction (DPI) of RNAi following the epitope-tagged target, Rieske protein (FS1) and CoxIV (CIV) by western blot. The parental cell line, in which LYRM7 is V5 tagged, but prior to transformation with inducible RNAi cassette (-) is also shown. Part of the stain-free gel is shown as a loading control. **(H)** Comparison of AlphaFold predicted structures of *T. brucei* LYRM7 (top) obtained from Wheeler (2021) and the *S. cerevisiae* LYMR7 ortholog (Google DeepMind dataset v6) known as Mzm1 (bottom). Keys to arrow heads on lower left and pLDDT score coloring on right.

**Figure S5. The inclusion of UQCRTBx1-4 yields a robust AlphaFold 3 prediction of the trypanosomal ubiquinol: cytochrome c oxidoreductase structure. (A)** Cord diagram mapping each binary interaction among the subunits integrated into the predicted Complex III structure, with either ≥0.7 (right diagram) or ≥0.9 (left) confidence scores. The number of contacts meeting these two confidence thresholds are listed on bottom left of each diagram. Each predicted polypeptide chain is depicted along the circumference of the circle as arc-shaped boxes colored by local pLDDT scores along their primary structure (N-to C-terminus, clockwise). Letter identifies each subunit and can be deciphered with Table S2. **(B)** Isolated view of the interaction interface connecting UQCRTBx2, UQCRTBx4,and Cytochrome C1 (CYC1). Inset expanded view of 2 and 1 non-covalent bonds among the subunits. Scale bar, 10 Å. **(C)** Alignment of predicted UQCRTBx1-4 UQCRTBx4, as integrated into the structure depicted in Figure 3D, aligned to the solved structure of *E. gracilis* ubiquinol: cytochrome c oxidoreductase (PDB 8IUF) (He *et al*., 2024). **(D)** Cord diagram as in A mapping of binary interactions within the predicted Complex III structure incorporating one copy of UQCRTBx2-V5. **(E)** Side-by-side comparison of ubiquinol: cytochrome c oxidoreductase incorporating 2 WT or one V5-tagged version of UQCRTBx2. Color and electrostatic charge keys on left and at bottom, respectively. Yellow dashed oval, wider opening; Teal dashed arrow, cytochrome c dock; arrowhead, V5 tag; *, location of missing UQCRB subunit in predicted structure.

**Figure. S6. Additional data regarding g-like proteins g’ and g’’. (A)** Multisequence alignment of *T. brucei* subunit g, g-like’ and g-like’’ with their orthologs from kinetoplastids listed on left. (**B)** Maximum likelihood tree of sequences aligned in A with subunit g subunits from the diplonemid (Dip) sister clade as an outgroup. Clade support >50 shown. Scale bar, number of substitutions per site. **(C)** SDS PAGE and western blot showing downregulation of the targeted protein in g’, g’’ and g’ & g’’ induced (+) RNAi cell lines versus uninduced negative controls (-). Proteins were detected via their epitope tags: g’’-V5 and g’-HA. Part of the stain-free gel is shown as a loading control**. (D)** Growth of PCF in which g’, g’’ and g’ & g’’ are depleted by inducible RNAi (RNAi+) in comparison to uninduced negative controls (RNAi-). Key in bottom plot. Standard deviation at each time point (n=3) not well visible at this scale. **(E)** BN PAGE and western blot detecting ATP synthase subunit β (ATPB) in cells induced for g’, g’’ and g’ & g’’ RNAi (+) versus uninduced controls (-) Total mitochondrial protein loaded shown at top. Positions of F1 subcomplex, ATP synthase monomer (F_O_F_1_) and the dimer ((F_O_F_1_)_2_) are marked on left.

**Figure S7. Comparison of complexomes from this study and Solayman and colleagues (2026). (A)** The numbers of total and mitochondrial proteins detected in all complexome studies. Individual datapoints are plotted on top of Tukey plots (grey outline). Symbol key, upper right (also for **E-G**); σ, standard deviation. **(B)** Total number of proteins detected in the indicated complexomes of PCF and BSF. Combine, our complexome datasets combining both BN PAGE gels; Solay., Solayman et al., (2026) (also for **C-H**). **(C)** Venn diagrams depicting intersecting and disjointed proteins detected in our complexome datasets (green, dataset indicated on left) and Solayman (blue). Stage indicated at the top. Numbers inside and outside circle intersection is the number of proteins shared in and exclusive to both studies, respectively. Brackets enclose the percentage of proteins from the study that are detected in the other dataset. Bottom row shows the average ± standard deviation of values above. U, our combined complexome. **(D)** Same as C, but for mitoproteome. **(E)** Portion of mitoproteome captured in the studies indicated on the bottom. Lines denote the average value in each column. *L. tar.*, denotes the symbol for the *L. tarentolae complexome* (used in **G**). (F) Portion of glycosomes-residing proteins (glycoproteome) captured in both studies. Labelling as in **E**. **(G)** Portion of mitoproteome captured in both studies organized according to the life cycle stage/species indicated on bottom. P, PCF; B, BSF. Labelling as in **E**. **(H)** The number of mitochondrial proteins detected in the datasets indicated as in **B**. Vacas denotes the number of proteins detected in isolated mitochondria from the indicated *T. brucei* life cycle stages (Vacas *et al*., 2026). **(I)** Size fraction distribution of proteins according to signal intensity as color coded (indicated on top right of upper graph across 60 fractions in Solayman et al., (2026) PCF (top) and BSF (bottom) complexomes. **(J)** t-SNE plot of *L. tarentolae* complexome (Solayman *et al*., 2026) with key proteins colored as indicated on top left.

**Figure S8. Comparison of GIP grouping of complexomes from this study and Solayman and colleagues (2026). (A)** The portion (% protein) of total protein assigned to a GIP group (GIP/Total) in the studies indicated below. Symbol key, right (also for **B**); σ, standard deviation. **(B)** The portion of the mitoproteome assigned to a GIP group (GIP/mito) as in **A**. **(C)** Venn diagram showing intersecting and disjoined proteins that were placed in a GIP group in both studies. As in **Fig. S7C. (D)** Presence of proteins indicated on left in GIP group indicated on top from both studies as indicated is demarked by a circle. These are compared to the proteins found in the solved Complex III structure in Solayman (Solay.) (2026), in green. True positive (+ve), black; false positive grey. **E.** Size fraction distribution of proteins according to normalized intensity as color coded top right of upper graph across 48 fractions in our PCF complexomes indicated on top left.

## Notes

### Competing Interest Statement

The authors have declared no competing interest.

### Summary of Updates

The manuscript has been updated according to the reviewers' comments.

