## Supplementary figures for "Interconnections with and within the trypanosomal respiratory chain revealed by complexome profiling"

Fig S1

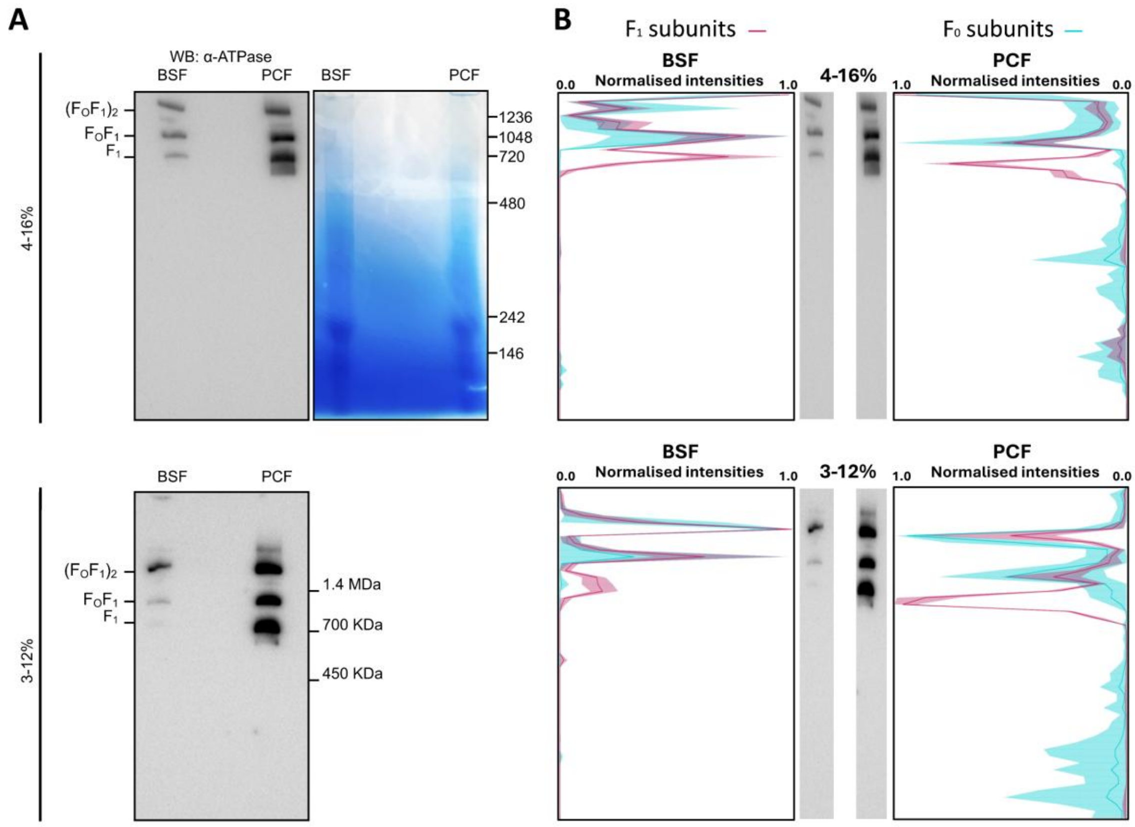

A
WB: α-ATPase
BSF
PCF
BSF
PCF
$(F_OF_1)_2$
$F_OF_1$
$F_1$
1236
1048
720
480
242
146
4-16%
BSF
PCF
1.4 MDa
700 KDa
450 KDa
3-12%
B
$F_1$ subunits
$F_0$ subunits
BSF
Normalised intensities
0.0
1.0
4-16%
PCF
Normalised intensities
1.0
0.0
BSF
Normalised intensities
0.0
1.0
3-12%
PCF
Normalised intensities
1.0
0.0

Fig S2

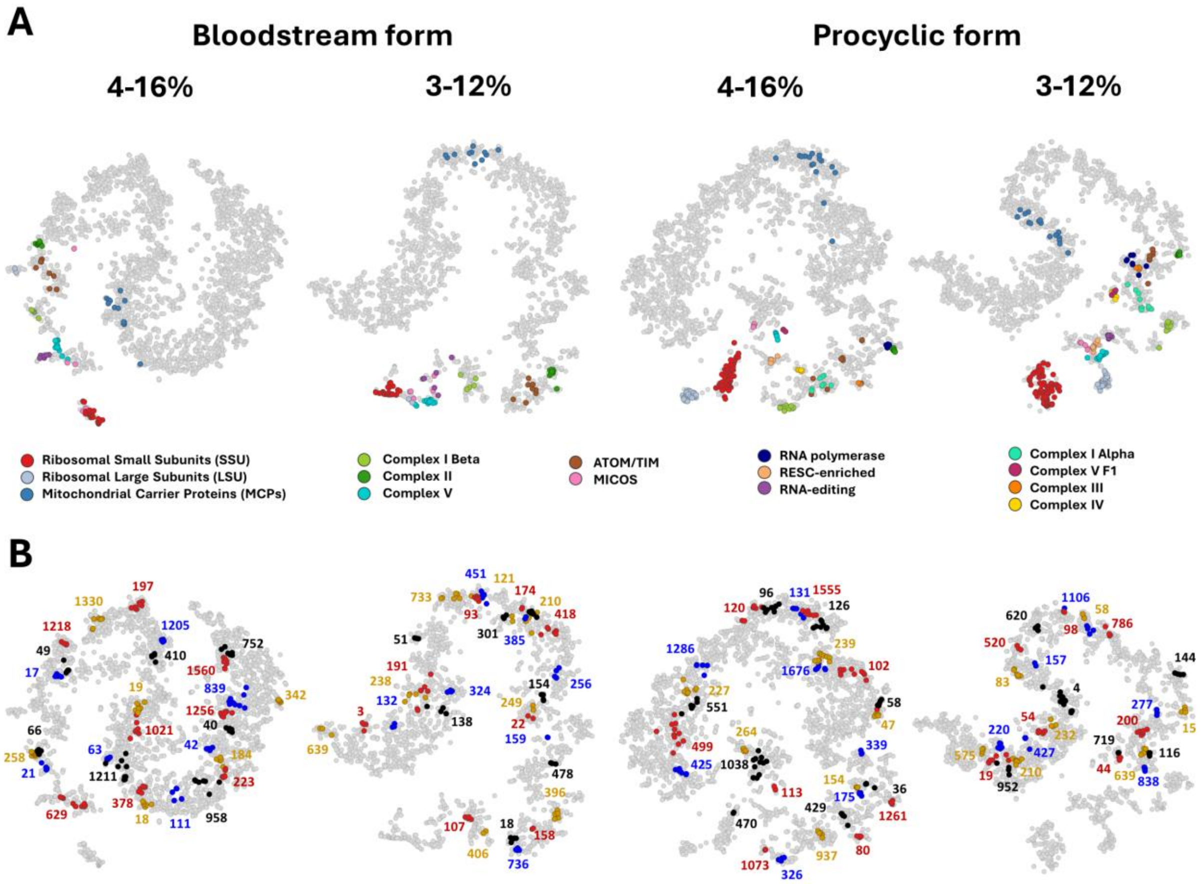
A
Bloodstream form
Procyclic form
4-16%
3-12%
4-16%
3-12%
Ribosomal Small Subunits (SSU)
Ribosomal Large Subunits (LSU)
Mitochondrial Carrier Proteins (MCPs)
Complex I Beta
Complex II
Complex V
ATOM/TIM
MICOS
RNA polymerase
RESC-enriched
RNA-editing
Complex I Alpha
Complex V F1
Complex III
Complex IV
B

Fig S3

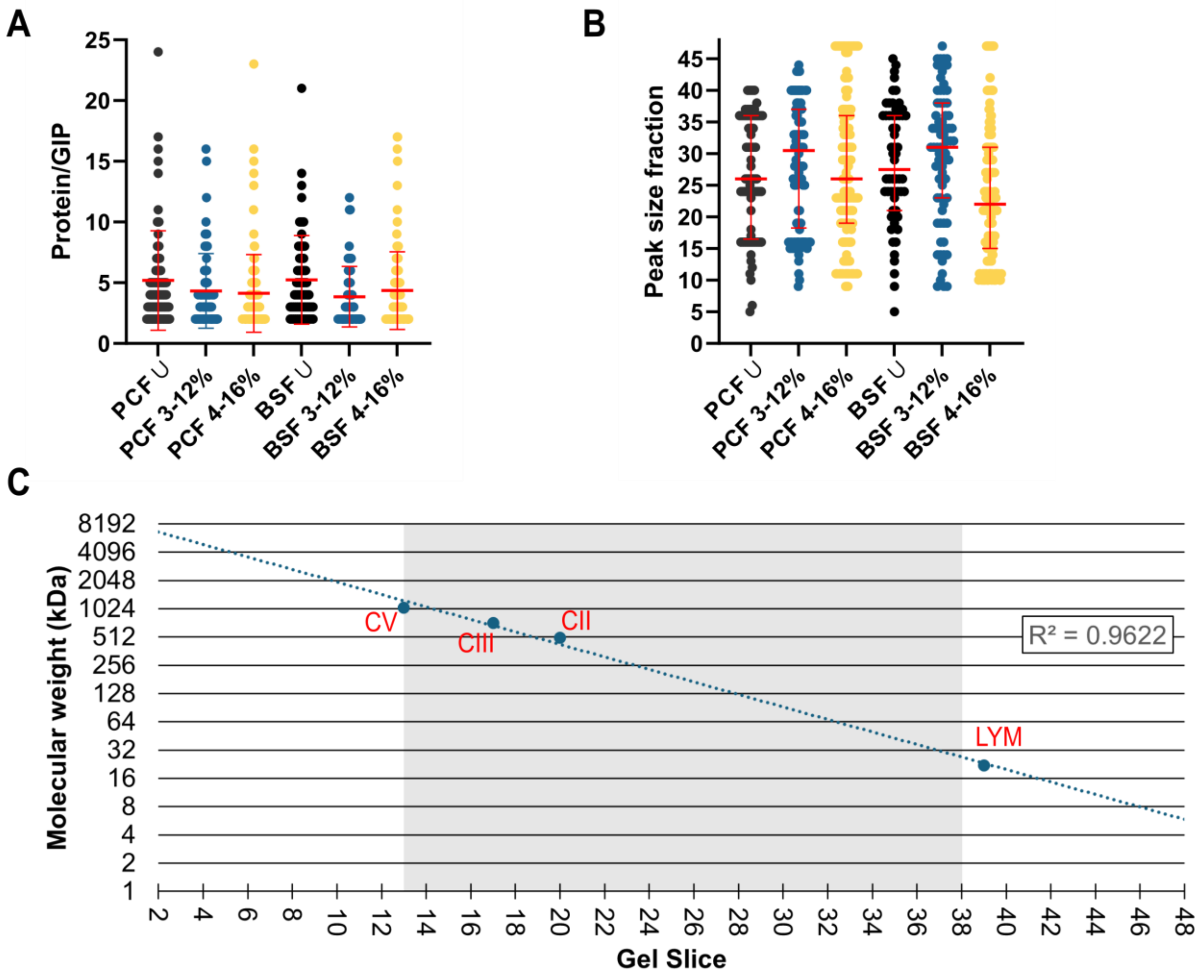
A
Protein/GIP
25
20
15
10
5
0
PCF U
PCF 3-12%
PCF 4-16%
BSF U
BSF 3-12%
BSF 4-16%
B
Peak size fraction
45
40
35
30
25
20
15
10
5
0
PCF U
PCF 3-12%
PCF 4-16%
BSF U
BSF 3-12%
BSF 4-16%
C
Molecular weight (kDa)
8192
4096
2048
1024
512
256
128
64
32
16
8
4
2
1
CV
CIII
CII
LYM
R² = 0.9622
Gel Slice
2 4 6 8 10 12 14 16 18 20 22 24 26 28 30 32 34 36 38 40 42 44 46 48

Fig S4

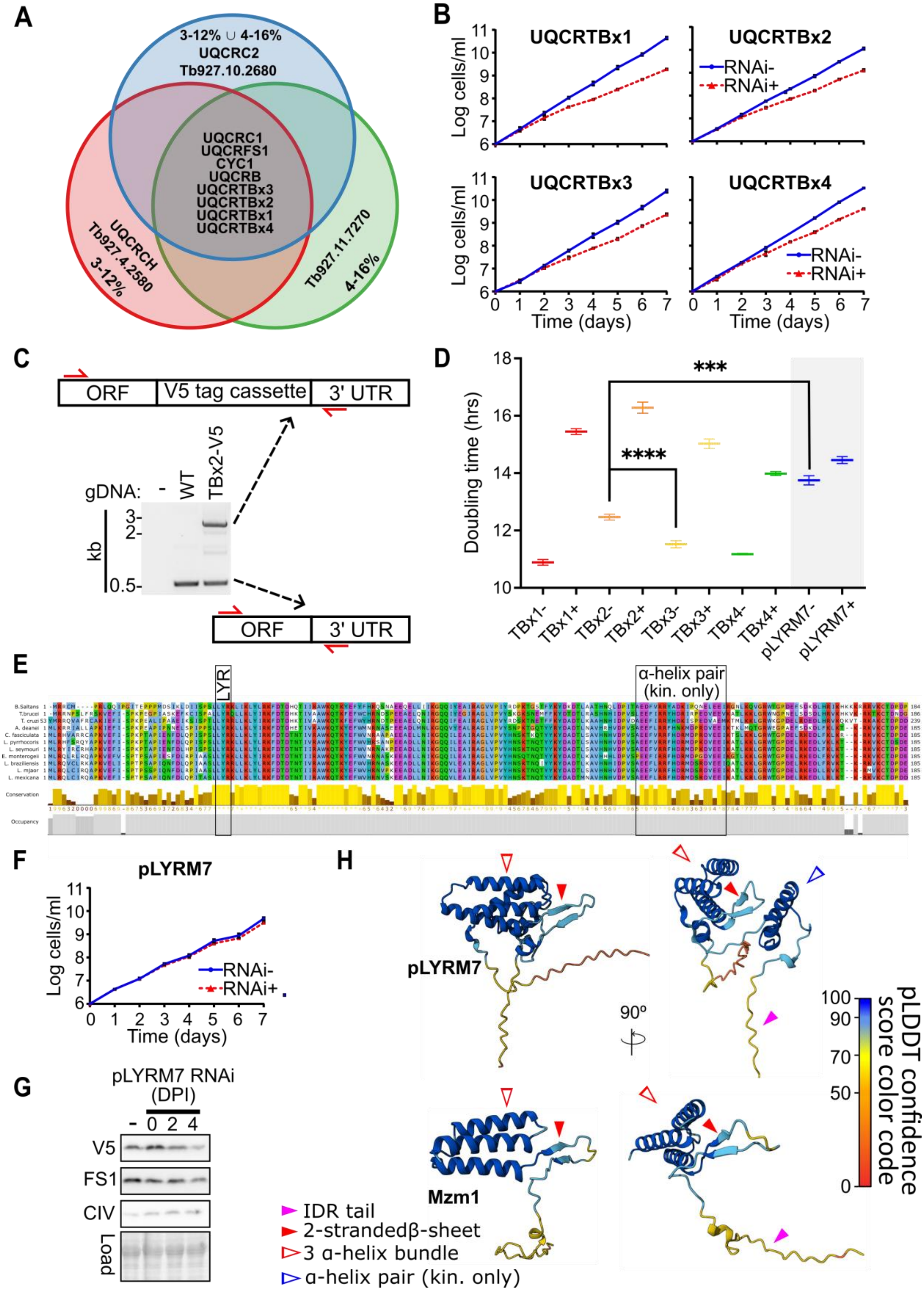

A
3-12% ∪ 4-16%
UQCRC2
Tb927.10.2680
UQCRC1
UQCRFS1
CYC1
UQCRB
UQCRTBx3
UQCRTBx2
UQCRTBx1
UQCRTBx4
UQCRCH
Tb927.4.2580
3-12%
Tb927.11.7270
4-16%
B
UQCRTBx1
UQCRTBx2
UQCRTBx3
UQCRTBx4
RNAi-
RNAi+
Log cells/ml
Time (days)
C
ORF
V5 tag cassette
3' UTR
gDNA:
-
WT
TBx2-V5
kb
3-
2-
0.5-
D
Doubling time (hrs)
***
****
TBx1-
TBx1+
TBx2-
TBx2+
TBx3-
TBx3+
TBx4-
TBx4+
pLYRM7-
pLYRM7+
E
LYR
α-helix pair
(kin. only)
Conservation
Occupancy
F
pLYRM7
Log cells/ml
Time (days)
RNAi-
RNAi+
G
pLYRM7 RNAi
(DPI)
-
0
2
4
V5
FS1
CIV
Load
H
pLYRM7
Mzm1
90º
IDR tail
2-strandedβ-sheet
3 α-helix bundle
α-helix pair (kin. only)
pLDDT confidence
score color code

Fig S5

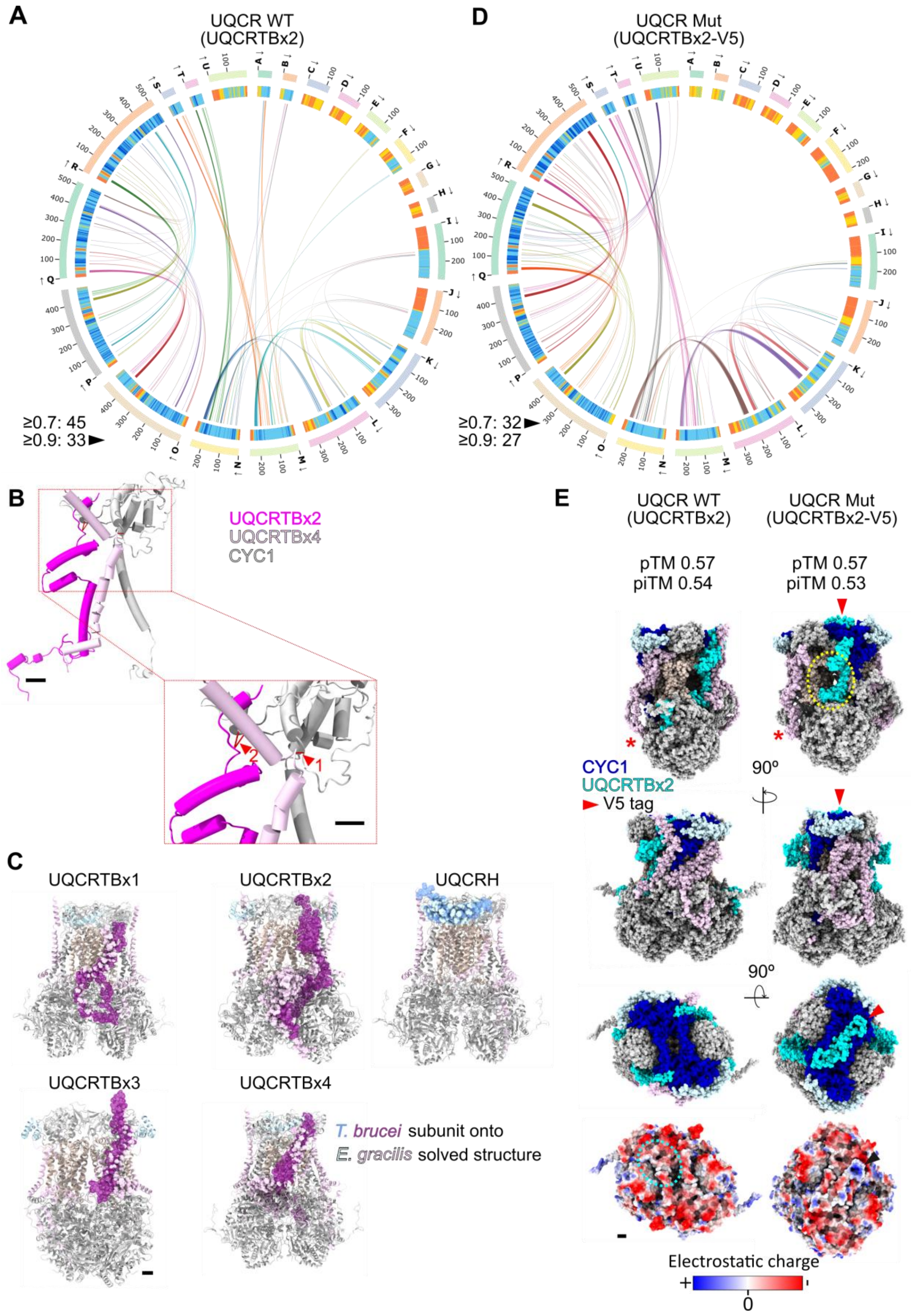

A
UQCR WT
(UQCRTBx2)
≥0.7: 45
≥0.9: 33
D
UQCR Mut
(UQCRTBx2-V5)
≥0.7: 32
≥0.9: 27
B
UQCRTBx2
UQCRTBx4
CYC1
C
UQCRTBx1
UQCRTBx2
UQCRH
UQCRTBx3
UQCRTBx4
T. brucei subunit onto
E. gracilis solved structure
E
UQCR WT
(UQCRTBx2)
UQCR Mut
(UQCRTBx2-V5)
pTM 0.57
piTM 0.54
pTM 0.57
piTM 0.53
CYC1
UQCRTBx2
V5 tag
90º
90º
Electrostatic charge
+
0
-

Fig S6

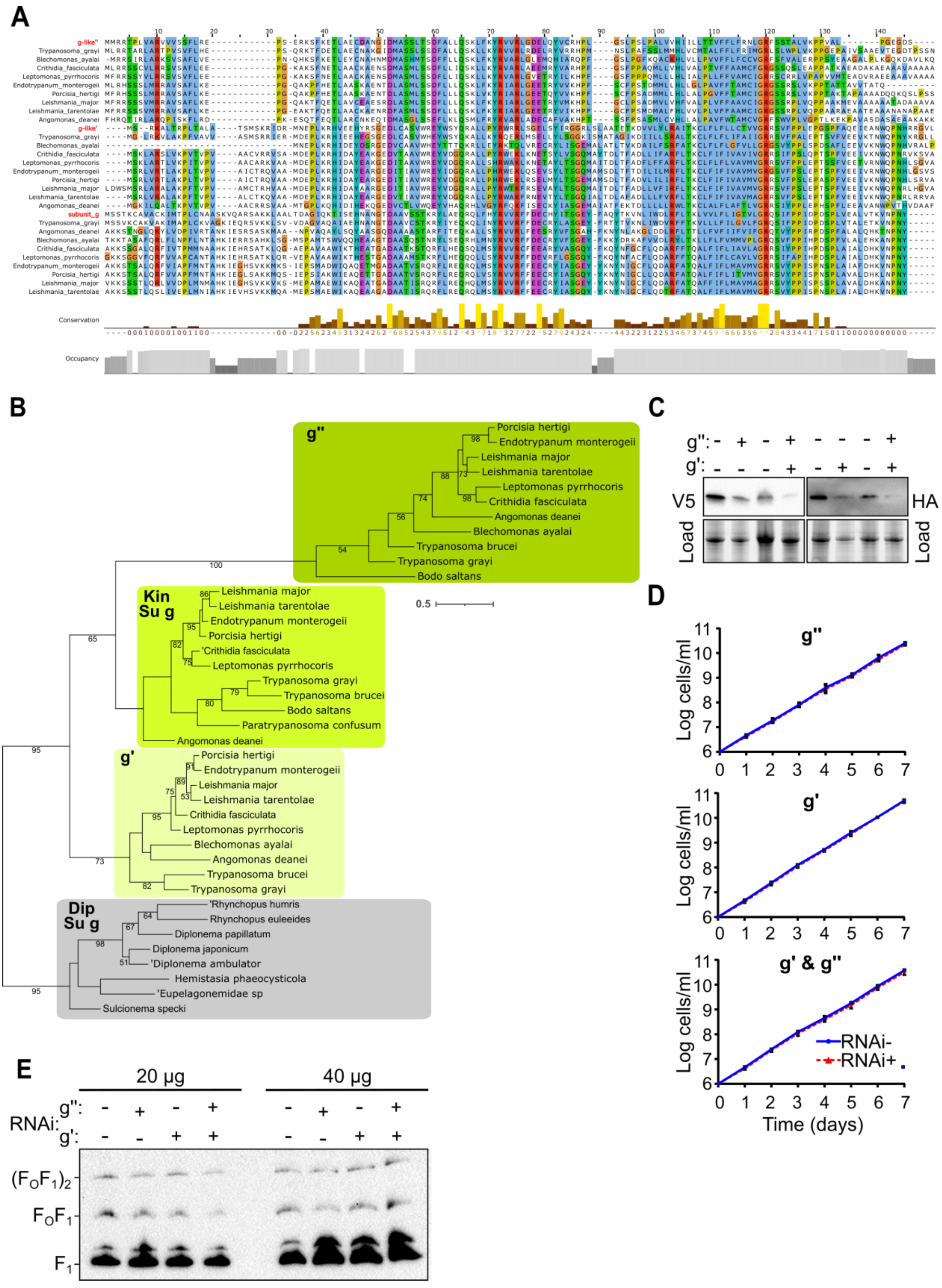

A
g-like''
Trypanosoma_grayi
Blechomonas_ayalai
Crithidia_fasciculata
Leptomonas_pyrrhocoris
Endotrypanum_monterogeii
Porcisia_hertigi
Leishmania_major
Leishmania_tarentolae
Angomonas_deanei
g-like'
subunit_g
Conservation
Occupancy
B
g''
Porcisia hertigi
Endotrypanum monterogeii
Leishmania major
Leishmania tarentolae
Leptomonas pyrrhocoris
Crithidia fasciculata
Angomonas deanei
Blechomonas ayalai
Trypanosoma brucei
Trypanosoma grayi
Bodo saltans
Kin Su g
Paratrypanosoma confusum
g'
Dip Su g
Rhynchopus humris
Rhynchopus euleeides
Diplonema papillatum
Diplonema japonicum
Diplonema ambulator
Hemistasia phaeocysticola
Eupelagonemidae sp
Sulcionema specki
0.5
C
g'':
g':
V5
HA
Load
D
g''
g'
g' & g''
Log cells/ml
Time (days)
RNAi-
RNAi+
E
20 µg
40 µg
RNAi:
g'':
g':
$(F_OF_1)_2$
$F_OF_1$
$F_1$

Fig S7

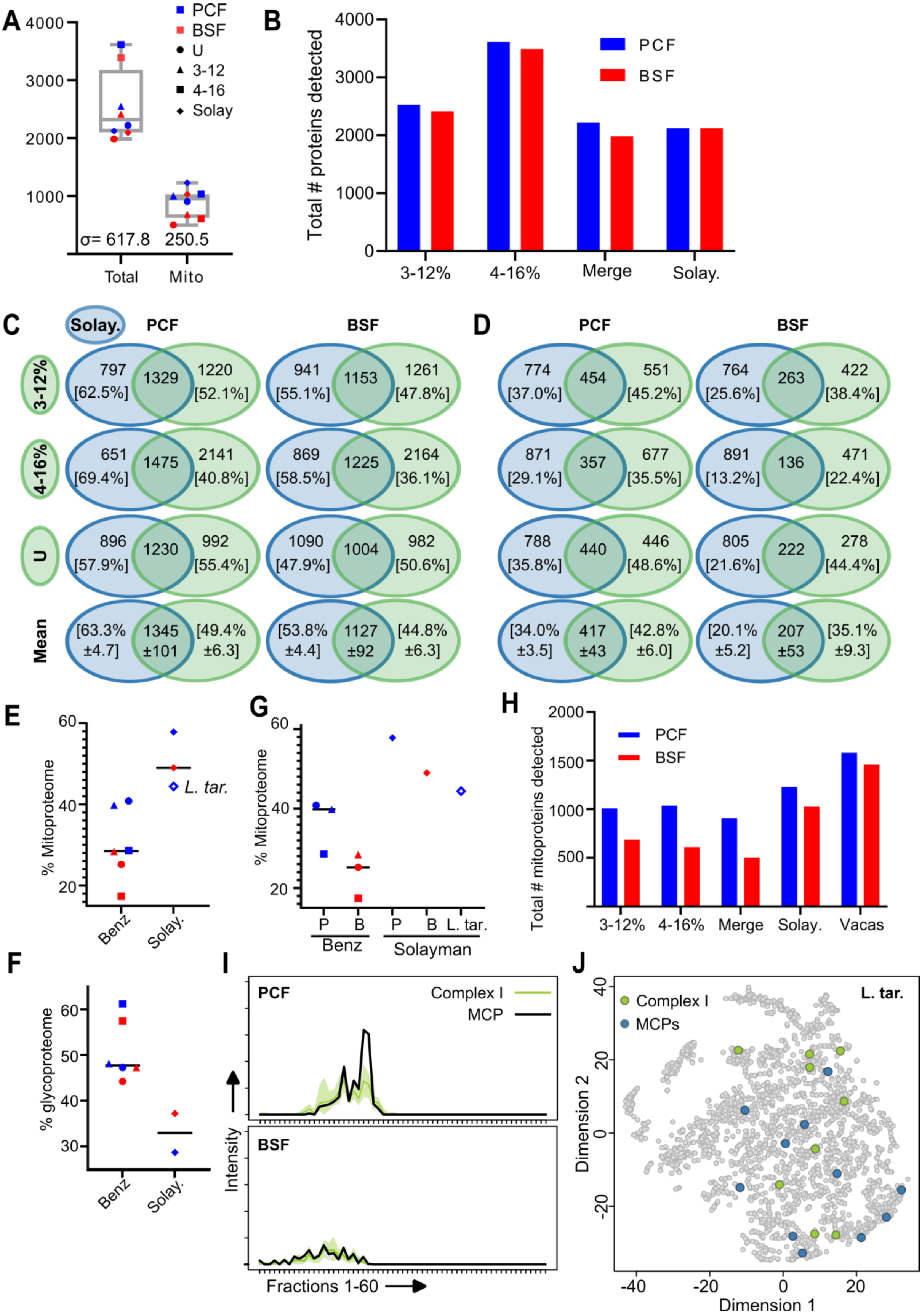
A
PCF
BSF
U
3-12
4-16
Solay
σ= 617.8
250.5
Total
Mito
B
Total # proteins detected
PCF
BSF
3-12%
4-16%
Merge
Solay.
C
Solay.
PCF
BSF
3-12%
4-16%
U
Mean
797 [62.5%] 1329 1220 [52.1%]
941 [55.1%] 1153 1261 [47.8%]
651 [69.4%] 1475 2141 [40.8%]
869 [58.5%] 1225 2164 [36.1%]
896 [57.9%] 1230 992 [55.4%]
1090 [47.9%] 1004 982 [50.6%]
[63.3% ±4.7] 1345 ±101 [49.4% ±6.3]
[53.8% ±4.4] 1127 ±92 [44.8% ±6.3]
D
PCF
BSF
774 [37.0%] 454 551 [45.2%]
764 [25.6%] 263 422 [38.4%]
871 [29.1%] 357 677 [35.5%]
891 [13.2%] 136 471 [22.4%]
788 [35.8%] 440 446 [48.6%]
805 [21.6%] 222 278 [44.4%]
[34.0% ±3.5] 417 ±43 [42.8% ±6.0]
[20.1% ±5.2] 207 ±53 [35.1% ±9.3]
E
% Mitoproteome
L. tar.
Benz
Solay.
G
% Mitoproteome
P
B
P
B
L. tar.
Benz
Solayman
H
Total # mitoproteins detected
PCF
BSF
3-12%
4-16%
Merge
Solay.
Vacas
F
% glycoproteome
Benz
Solay.
I
PCF
Complex I
MCP
BSF
Intensity
Fractions 1-60
J
Complex I
MCPs
L. tar.
Dimension 2
Dimension 1

Fig S8

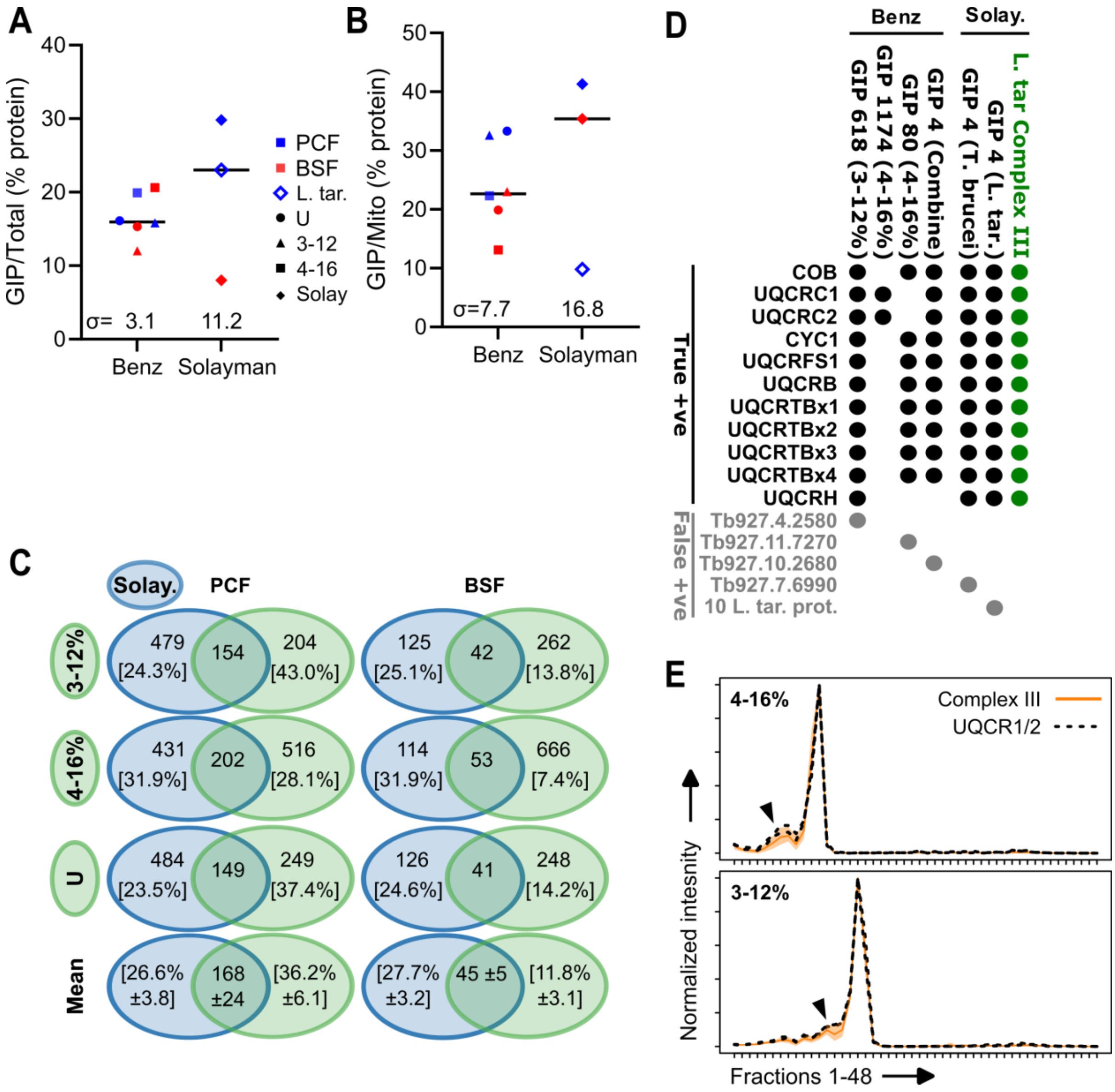

A
GIP/Total (% protein)
σ= 3.1
11.2
Benz
Solayman
PCF
BSF
L. tar.
U
3-12
4-16
Solay
B
GIP/Mito (% protein)
σ=7.7
16.8
Benz
Solayman
C
Solay.
PCF
BSF
3-12%
479 [24.3%]
154
204 [43.0%]
125 [25.1%]
42
262 [13.8%]
4-16%
431 [31.9%]
202
516 [28.1%]
114 [31.9%]
53
666 [7.4%]
U
484 [23.5%]
149
249 [37.4%]
126 [24.6%]
41
248 [14.2%]
Mean
[26.6% ±3.8]
168 ±24
[36.2% ±6.1]
[27.7% ±3.2]
45 ±5
[11.8% ±3.1]
D
Benz
Solay.
GIP 618 (3-12%)
GIP 1174 (4-16%)
GIP 80 (4-16%)
GIP 4 (Combine)
GIP 4 (T. brucei)
GIP 4 (L. tar.)
L. tar Complex III
True +ve
COB
UQCRC1
UQCRC2
CYC1
UQCRFS1
UQCRB
UQCRTBx1
UQCRTBx2
UQCRTBx3
UQCRTBx4
UQCRH
False +ve
Tb927.4.2580
Tb927.11.7270
Tb927.10.2680
Tb927.7.6990
10 L. tar. prot.
E
4-16%
3-12%
Complex III
UQCR1/2
Normalized intesnity
Fractions 1-48
